# Structural basis of SIRT7 nucleosome engagement and substrate specificity

**DOI:** 10.1101/2024.10.10.617549

**Authors:** Carlos Moreno-Yruela, Babatunde E. Ekundayo, Polina N. Foteva, Esther Calvino-Sanles, Dongchun Ni, Henning Stahlberg, Beat Fierz

## Abstract

Chromatin-modifying enzymes selectively target distinct residues within histones to finetune gene expression profiles. SIRT7 is an NAD^+^-dependent histone deacylase often deregulated in cancer, which deacetylates either H3 lysine 36 (H3K36) or H3K18 with high specificity within nucleosomes. Here, we report structures of nucleosome-bound SIRT7, and uncover the structural basis of its specificity towards H3K36 and K18 deacylation, combining a mechanism-based cross-linking strategy, cryo-EM, mutagenesis and enzymatic assays. We show that the SIRT7 N-terminus represents a unique, extended nucleosome-binding domain, reaching across the nucleosomal surface to the acidic patch. The catalytic domain binds at the H3-tail exit site, engaging both DNA gyres of the nucleosome. Contacting H3K36 versus H3K18 requires a change in enzyme binding pose, and results in structural changes in both SIRT7 and the nucleosome. These structures reveal interactions critical for target lysine specificity, allowing us to engineer enzyme activity towards H3K18 or 36, and provides a basis for small molecule modulator development.

---

Sirtuins are ubiquitous regulators of protein function through NAD^+^-dependent cleavage of *N*^ε^-acyl-lysine post-translational modifications (PTMs)^1,2^, and deeply involved in chromatin regulation via histone deacetylation^3^. The sirtuin family in mammals consists of seven members corresponding to class I (SIRT1‒3), class II (SIRT4), class III (SIRT5), and class IV (SIRT6, SIRT7)^4^. Four of these enzymes ‒ SIRT1, SIRT2, SIRT6 and SIRT7 ‒ erase histone lysine acetylation (Kac) to regulate chromatin structure and gene transcription, and they have isozyme-specific roles as epigenetic regulators of disease pathways in cancer, neurodegeneration, and cardiometabolic disorders^3,5,6^.

SIRT7 is a nuclear enzyme both present in the nuclear lumen and the nucleolus. In the latter, it controls ribosomal DNA transcription^7,8^ and maintains chromatin packing as a protection from homologous recombination and damage^9,10^. Outside of the nucleolus, SIRT7 is responsible for gene promoter regulation and especially relevant in cancer, where it is often overexpressed and helps to repress tumour suppressor genes and activate metastatic pathways^11–15^. These functions depend on SIRT7-mediated deacetylation of nucleolar proteins and of the histone mark H3K18ac^11,16,17^. Accordingly, SIRT7 inhibitors have been proposed as potential epigenetic cancer chemotherapy^18^.

SIRT7 is also a key regulator of H3K36ac in cells^15,19^. This modification lowers nucleosome stability^19^ and favours euchromatin formation^20^, and it is enriched at transcription start sites^20^. Importantly, H3K36ac competes with methylation of H3K36, a PTM that plays a role in many pathways including DNA damage-signalling^21^, which may explain the role of SIRT7 in DNA repair^19^. Together, the H3K18 and H3K36 deacetylase activities of SIRT7 render it an important regulator of gene expression and chromatin integrity. However, little is known about the molecular mechanisms governing SIRT7 function, and recent studies underscore that its role in cancer is poorly understood and highly context dependent, leading to both tumour promotion and tumour suppression^22–24^.

SIRT7 activity shows characteristics unlike those of any other sirtuin. *In vitro*, SIRT7 presents both lysine deacylase and ADP-ribosylase activities^11,25–27^, and it has preference for the long-chain acyl-lysine modification decanoyl-lysine (Kdec)^28^. Its deacylase activity on peptide and protein substrates requires binding to nucleic acids, in contrast with the rest of the sirtuin family^29,30^. SIRT7 is most active when bound to nucleosomes, showing affinity in the nanomolar range towards unmodified nucleosomes^31^ and targeting H3K18ac and H3K36ac nucleosome substrates with unique selectivity^19,32,33^. The main structural difference of SIRT7, which may explain its behaviour, are the two positively-charged domains that flank its catalytic domain, proposed to mediate nucleic acid binding and activation^29,30,34,35^. A crystal structure of a part of the N-terminal domain shows helical folding reminiscent of DNA-binding domains^36^, but no other structural evidence has been provided to date.

Here, we reveal the structural basis for SIRT7 specific substrate recognition and activation by nucleosomes. We synthesised nucleosomes with mechanism-based inhibitors at lysines 36 and 18 in H3 to trap the active conformation of SIRT7^37,38^. We then solved the structures of SIRT7 bound to a nucleosome and contacting H3K36 or H3K18 by cryogenic electron microscopy (cryo-EM) to a resolution of 2.8 and 3.5 Å respectively, revealing its atypical nucleosome binding mode: we found that SIRT7 is recruited to the nucleosome by its extended N-terminal domain, which traverses the nucleosome surface and binds to both the acidic patch and nucleosomal DNA. At the same time, the catalytic domain engages both DNA gyres at the H3-tail exit site of the nucleosome. Strikingly, SIRT7 adopts different conformations depending on the target lysine, H3K36 or H3K18. Moreover, by comparing H3K18- and H3K36-bound structures, we found that distortion of the nucleosomal DNA structure is only needed for H3K18 targeting. Finally, structure-based mutagenesis afforded SIRT7 variants with altered substrate selectivity and unveiled a key catalytic domain loop responsible for the H3K36-specific targeting of SIRT7.

## Results

### Thiourea histone modifications afford mechanism-based SIRT7:nucleosome complexes

SIRT7 is highly activated by nucleosome binding and is most active against H3K36 substrates, followed by H3K18^19,31^. To reveal the structural basis of how this enzyme can specifically target two distinct sites within a nucleosome, we decided to stabilise nucleosome-bound SIRT7 contacting H3K36 or H3K18. We envisioned the use of mechanism-based sirtuin inhibitor warheads at these two positions. Typical functional groups are thioureas^38,39^, which substitute nicotinamide at the 1’ ribose position of NAD^+^ and stall the sirtuin mechanism at one of the covalent intermediates^37,40^, or form an alternative iminium adduct, as recently found in a SIRT6:thiourea complex (Fig. 1A)^38^. To generate these constructs, we first introduced methylthiourea (MTU) and decylthiourea (DTU) modifications onto synthetic H3 peptides on resin^41^. Then, we followed with two or three fragment native chemical ligation (NCL) protocols to obtain full-length H3 modified at K18 or K36, respectively (Fig. 1B)^42^.

**Fig. 1.**
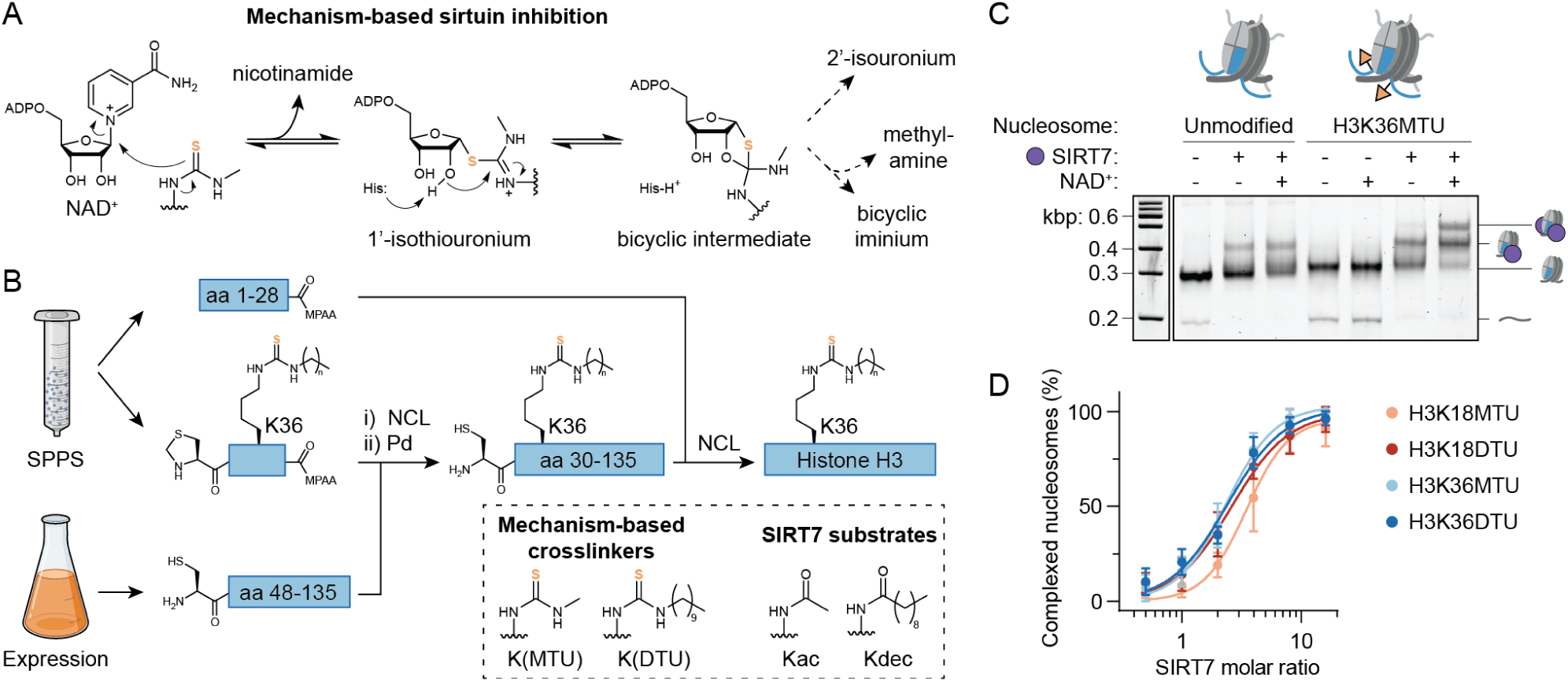
Thiourea-modified histone H3 stabilizes the SIRT7-nucleosome interaction. **A**, Mechanism-based sirtuin inhibition by thioureas^37,38^. **B**, Semi-synthetic approach to generate full-length H3 with thiourea or amide modifications at K36 (see Supporting Information for K18 modifications). Peptides covering amino acids (aa) 1‒ 28 and 29‒46 were prepared by solid phase peptide synthesis (SPPS) and linked to a recombinant fragment by NCL. MPAA: 4-mercaptophenylacetic acid. Overall H3 yield: 12‒22% in 0.5‒1.1 mg scale. See Supplementary Fig. 1A for sample HPLC and MS spectra. **C**, SIRT7 EMSA of nucleosome samples with and without H3K36MTU modification (SIRT7-nucleosome molar ratio: 3). See Supplementary Fig. 1B for H3K18DTU EMSA. **D**, Quantification of SIRT7-nucleosome complex formation with H3K18 or H3K36 thioureas. Error bars represent mean ± SD (*n* = 3). See Supplementary Fig. 1C for representative EMSAs.

With the full-length H3 constructs in hand, we reconstituted nucleosomes using the nucleosome-positioning 601 DNA sequence^43^ with 20 bp flanking additions that promote SIRT7 activity at H3K36ac^19^. Thiourea-containing nucleosomes formed more stable complexes with SIRT7 than unmodified nucleosomes, as observed by electrophoretic mobility shift assay (EMSA). These complexes followed 1:1 as well as 2:1 stoichiometry, where both H3 molecules are engaged by SIRT7 (Fig. 1C and Supplementary Fig. 1B). Importantly, complex stabilization was NAD^+^-dependent, which demonstrated the formation of mechanism-based adducts at the thiourea position (Fig. 1A). Complexes linked at H3K36 and at H3K18 were stabilised with similar performance (Fig. 1D and Supplementary Fig. 1C), with both the shorter MTU and longer DTU functionalities that mimic the preferred Kac and Kdec substrates of SIRT7 (Fig. 1B)^28,44^. Thus, mechanism-based adduct formation does not depend on relative enzyme specificity.

### SIRT7 binds the nucleosome acidic patch and both DNA gyres to target H3K36

We proceeded to analyze the complex of full-length SIRT7 with H3K36MTU-containing nucleosomes by single-particle cryo-EM (Supplementary Fig. 2A). Three-dimensional (3D) reconstruction to an overall resolution of 2.8 Å displayed SIRT7 bound to the nucleosome edge (Fig. 2A and Supplementary Fig. 3 and 4). The cryo-EM map revealed distinct densities for SIRT7 side chains, especially at the nucleosome-binding interface and for the mechanism-based adduct on the nucleosome, which enabled the detailed modelling of a high-resolution structure of the complex (Fig. 2B and Supplementary Fig. 5).

**Fig. 2.**
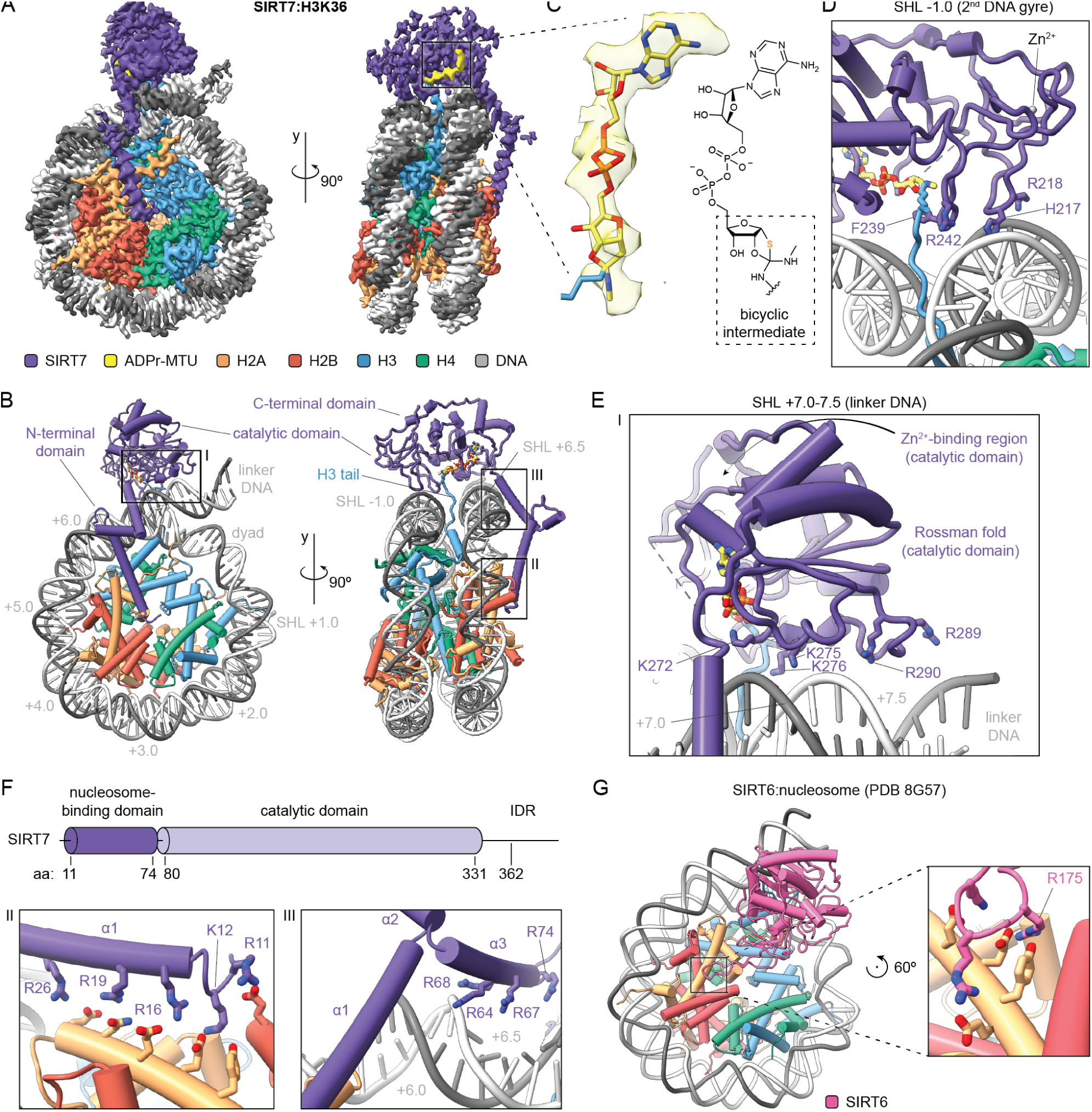
Cryo-EM structure of SIRT7 bound to the H3K36MTU nucleosome. **A**, Cryo-EM reconstruction showing SIRT7 bound to the nucleosome, with a mechanism-based cross-link to H3K36. See Supplementary Figs. 3‒4 for data processing workflow, resolution and angular particle distribution. **B**, Model of SIRT7(10-362) bound to the H3K36MTU nucleosome. See Supplementary Fig. 5 for model quality. The grey numbers indicate superhelical locations (SHL) along the nucleosomal DNA. **C**, Cryo-EM density of the ADPr-MTU covalent adduct and fitting of a mechanism-based bicyclic intermediate model. For a detailed view on regions I-III, see panels E and F. **D**, Detailed interactions of the SIRT7 catalytic domain with the 2^nd^ DNA gyre and the H3 tail. **E**, Detailed interactions with linker DNA, corresponding to frame “I” of the model as shown in Fig. 2B. **F**, SIRT7 domain organization and detailed interactions of the N-terminal (nucleosome binding-) domain with DNA and histones. Panels correspond to “II” and “III” portions of the model. **G**, Cryo-EM structure of SIRT6 bound to a nucleosome, highlighting the arginine anchor (R175) interaction with the nucleosome acidic patch (PDB 8G57)^46^.

In our model, the catalytic domain binds to the side of the nucleosome at the H3 tail exit site. The SIRT7 substrate pocket accommodates the H3K36 side chain securing it in place using residue F239. The map permitted reconstruction of the mechanism-based adenosine diphosphate ribose (ADPr)-MTU conjugate (Fig. 2C), consistent with the bicyclic intermediate also observed in a X-ray structure of SIRT5 bound to a thioamide^45^. This was in contrast with a recent structure where ADPr in SIRT6 formed an iminium link with H3K9MTU due to elimination of methylamine^38^ (Fig. 1C). Here, our cryo-EM density clearly indicated that the methylamine substituent is still present, and it further validated mechanism-based complex formation.

Contrary to enzymes that methylate^47^ or demethylate^48^ H3K36, SIRT7 does not unwrap DNA to target this residue from the histone octamer face. Instead, the SIRT7 catalytic domain presents two basic patches that bind in a cross-DNA gyre conformation (Fig. 2B), similar to PWWP H3K36me3 reader domains (Supplementary Fig. 6)^49^. In this configuration, multiple loops surrounding the SIRT7 catalytic pocket establish contacts with the backbone of both gyres of nucleosomal DNA as well as with extra-nucleosomal ‘linker’ DNA. Three residues away from F239, the side chain of R242 binds DNA (3.1 Å distance) at the 2^nd^ DNA gyre (superhelical location, SHL, -1.0). Here the DNA is also contacted by a neighbouring loop within the Zn^2+^-binding region of SIRT7 via residues H217 and R218 (3.6‒4.6 Å, Fig. 2D). Opposite of F239, a highly conserved SIRT7 loop^35^ within the NAD^+^-binding Rossmann fold makes weak electrostatic interactions (4.7‒6.2 Å) with DNA at SHL +7.0, using lysines 272, 275 and 276 (Fig. 2E). The Rossman fold further extends along the linker DNA and reaches the phosphate backbone at SHL +7.5 with R289 (3.8 Å) and, to a lesser extent, R290 (6.2 Å).

Remarkably, SIRT7 interacts with the octamer face and the H2A‒H2B acidic patch via its N-terminal domain, which folds into three α-helices that extend across the nucleosome (Fig. 2B). Helix α1 binds the classical acidic patch residues E105 of H2B with R11, and E61, D90 and E92 of H2A with K12 (Fig. 2F). The SIRT7 α1 helix then follows the H2A helix and further contacts H2A E64, N68 and D72 with arginines 16, 19 and 26 (Fig. 2F), and it forms weak electrostatic interactions (5.4‒6.1 Å) with arginines 18 and 21. The end of helix α1 sits close to SHL +6.0 and features three additional arginines, 30, 34 and 37, of which only R37 makes a direct contact (4.9 Å) with the DNA phosphate backbone. Helix α2 serves as the ‘elbow’ of the N-terminal domain and lets helix α3 accommodate in the major groove at SHL +6.5, where it contacts DNA with arginines 64, 68 and 74 (2.8‒3.3 Å, Fig. 2F). Thus, the SIRT7 N-terminal domain serves as a multivalent nucleosome-binding domain through eight side chains forming strong electrostatic interactions, among other contacts. This atypical binding mode explains how SIRT7 can target preferentially H3K36ac PTMs on the side of the nucleosome while interacting with the acidic patch^31^.

Multiple nucleosome-modifying enzymes interact with the acidic patch for chromatin binding, often through a single arginine anchor^50,51^. The closely related sirtuin SIRT6 shares such binding mode thanks to a specific loop insertion within the catalytic domain^38,46^ (Fig. 2G). In contrast, according to our structure the SIRT7 catalytic domain sits much further from the centre of the nucleosome, and the N-terminal domain interacts with the same anchor hotspot via an extended set of basic residues from R11 to R16, exhibiting non-canonical acidic patch interactions unlike any other enzyme shown to date (Fig. 2F)^51^.

Finally, three limited regions are missing in our SIRT7 model: the outermost N-terminus (aa 1‒9), a flexible loop within the catalytic domain (aa 120‒138), and part of the C-terminal domain (aa 363‒400). The loop around aa 120-138 is predicted by AlphaFold 3^52^ to contact DNA close to the nucleosome-binding domain (Supplementary Fig. 7), albeit at lower confidence than the rest of the catalytic domain. In agreement to this prediction, our cryo-EM map shows low-resolution density within the corresponding region, which suggests that this loop is disordered and may contribute to nucleosome binding. As for the C-terminal tail, this fragment is predicted to be an intrinsically-disordered region (IDR, Fig. 2F) containing a nuclear-localization sequence^35^. The C-terminal tail is also predicted to bind nucleic acids and has been shown to mediate activation by DNA^29^. In our model, a part of it is visible contacting the top of the catalytic domain (Fig. 2B). The remaining IDR may further project into the solution, allowing to form nucleic acid interactions in the vicinity.

### Targeting H3K18 involves structural changes in both SIRT7 and the nucleosome

To evaluate how SIRT7 contacts its substrate at H3K18 and to compare target-lysine specific binding modes, we next reconstituted a SIRT7 complex with a H3K18DTU-modified nucleosome for analysis by cryo-EM (Supplementary Fig. 2B). However, while the mechanism-based crosslink performed equally well as compared to the H3K36MTU (Fig. 1D), the SIRT7:H3K18DTU nucleosome complex required further stabilisation by gradient fixation^53^ in order to obtain density in the cryo-EM maps. This suggests that SIRT7 binding is more dynamic when bound to H3K18 than to H3K36. Nonetheless, we obtained a cryo-EM map of the SIRT7: H3K18DTU complex at 3.5 Å resolution overall and 6 Å on average for SIRT7 (Fig. 3A and Supplementary Figs. 8‒9), which enabled modelling of the SIRT7 domains into the cryo-EM map (Fig. 3B and Supplementary Fig. 10A).

**Fig. 3.**
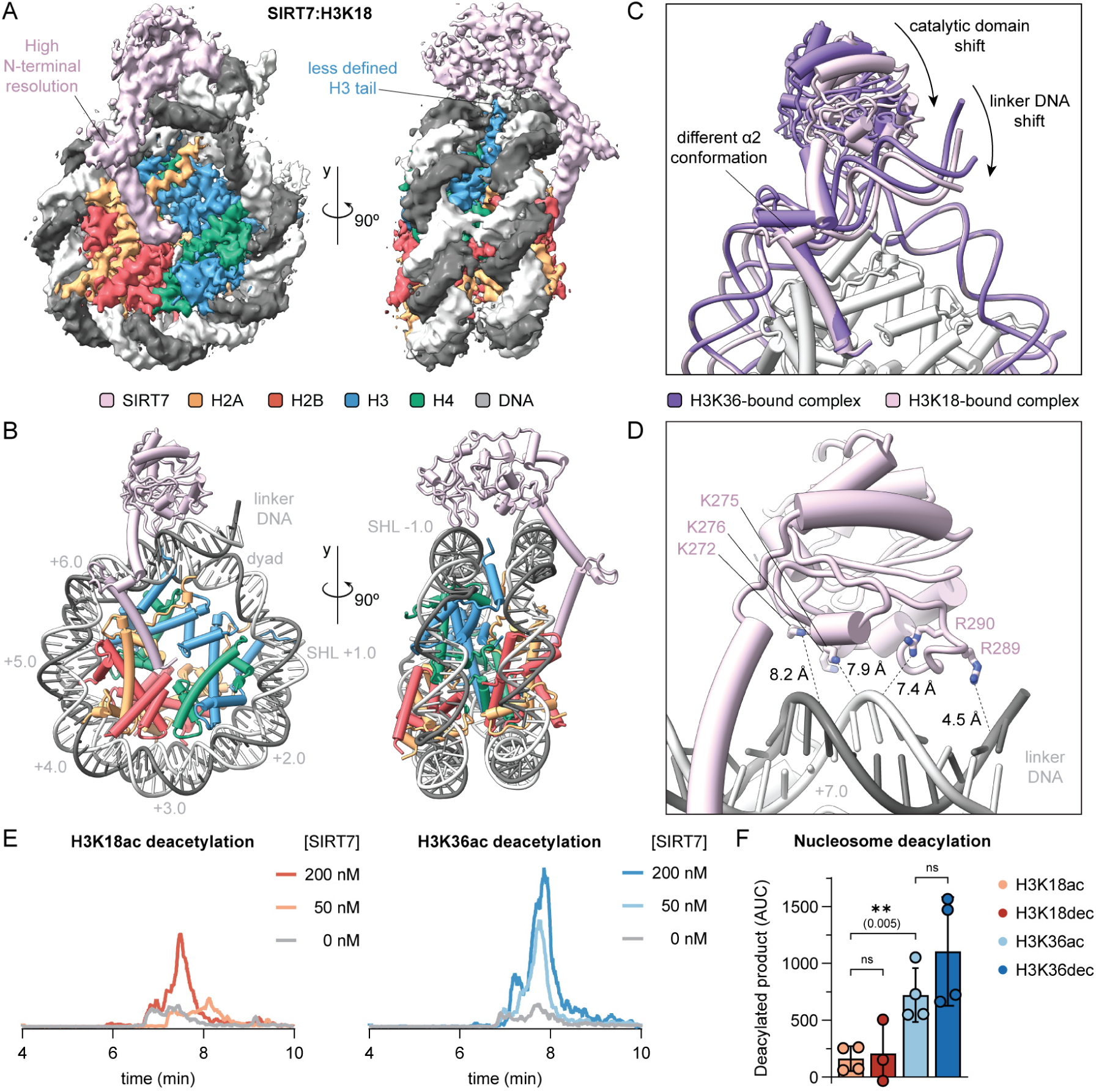
Altered binding pose of SIRT7 in complex with the H3K18DTU nucleosome. **A**, Cryo-EM reconstruction showing SIRT7 bound to the H3K18DTU nucleosome. See Supplementary Figs. 8‒9 for data processing workflow, resolution and angular particle distribution. **B**, Model of SIRT7(10-362) bound to the H3K18DTU nucleosome. See Supplementary Fig. 10 for model quality. **C**, Comparison of SIRT7:nucleosome complexes bound to the H3K36 and H3K18 positions, with highlighted key differences. **D**, Distance between residues that contact DNA for H3K36 binding and the DNA backbone in the H3K18-bound conformation. **E**, LC-MS analysis of nucleosome deacetylation by SIRT7, showing the extracted ion chromatogram of the corresponding deacetylated product ([M+21H]^21+^ ion). Nucleosome concentration was 200 nM. **F**, LC-MS quantification of nucleosome deacylation by SIRT7 (50 nM concentration). Data represent the area under the curve (AUC, [M+21H]^21+^ ion) of extracted ion chromatograms minus background. Error bars represent mean ± SD (*n* ≥ 3), ns: p > 0.05.

The N-terminal and catalytic domains of SIRT7 interact with similar nucleosome regions in both the H3K18-bound and H3K36-bound models, though they exhibit key differences in orientation that emphasise a substrate-specific conformation (Fig. 3C). Notably, the catalytic domain adopts a distinct conformation when bound to H3K18, indicating that movement within this domain is necessary for H3K18 targeting (Fig. 3C).

In concert with this reorientation of SIRT7, the linker DNA contacting the catalytic domain is bent inwards towards the face of the nucleosome by 6° relative to its position in the H3K36-bound structure (Supplementary Fig. 10B). This shift of the linker DNA allows a contact with the H2A C-terminal tail that has also already been observed in histone H1-induced linker DNA bending (Supplementary Fig. 10C)^54–56^. Moreover, R289 forms a closer interaction with the linker DNA region, whereas other residues contacting both linker DNA (K272, 275 and 276, and R290) and nucleosomal DNA (H217, R218 and R242), are positioned further away from the DNA backbone (>7 Å, Fig. 3D). These changes in distance reveal weakened electrostatic interactions at the SIRT7-DNA interface when targeting H3K18, possibly accounting for the less defined cryo-EM density and indicating increased flexibility compared to the H3K36-bound map.

Together with the overall reduced resolution of the SIRT7 catalytic domain in comparison to the rest of the cryo-EM map, we found less defined density in the substrate-binding pocket. In consequence we did not model the H3 tail or the ADPr-DTU crosslink (Fig. 3A). Since the distance between the substrate-locking F239 residue and the H3 tail exit site is similar in this model compared to the H3K36-bound structure, the H3 tail must form a loop along the linker

DNA to place the H3K18 side chain in the SIRT7 active pocket. In agreement, we observed additional putative H3 density within the open space generated by the 6° displacement of the linker DNA (Supplementary Fig. 10C).

The shift of the catalytic domain further alters the overall conformation of the nucleosome-binding domain and in particular changes the bending angle of helix α2 by 30° (Fig. 3C), which in this case is more similar to the published partial crystal structure of the SIRT7 N-terminus^36^. Importantly, both helices α1 and α3 maintain contacts with the acidic patch and the DNA compared to the previous conformation, which suggests that the positioning of the nucleosome-binding domain is primarily mediated by these key interactions and not by the location of the PTM on the nucleosome. In contrast, the hinge formed around helix α2 generates the necessary flexibility for the enzyme to shift pose.

The overall comparison of the two cryo-EM datasets indicates that SIRT7 adopts a more defined conformation for H3K36 than for H3K18 binding, the latter of which also includes a distortion of nucleosomal DNA conformation. To test how these different binding modes relate to enzymatic activity of SIRT7, we performed deacetylation assays. LC-MS analysis of the products revealed higher SIRT7 activity on H3K36ac compared to H3K18ac substrates (>4-fold higher activity at 50 nM SIRT7 concentration, Fig. 3C), in line with previous reports^19,32^. Furthermore, although SIRT7 shows greater activity towards long acyl modifications in peptide substrates^28^, we did not observe significant differences between Kac and Kdec substrates at either H3K36 or H3K18 within nucleosomes (Fig. 3D). In summary, our results indicate that nucleosome binding directs SIRT7 activity towards deacetylation of H3K36.

### The nucleosome-binding domain recruits SIRT7 to chromatin through multivalent interactions

Using the two structures for H3K36 and H3K18-contacting SIRT7 as a basis, we investigated how the observed interactions of the N-terminal nucleosome-binding domain with the histone octamer and in particular the acidic patch drive overall binding and activity. To this end, we generated two new SIRT7 variants. In the first, we introduced two mutations into the nucleosome-binding domain, targeting the arginine anchor that directly contacts the acidic patch (SIRT7 R11A, K12A). In a second variant, we removed the complete helix α1 of the nucleosome-binding domain (SIRT7 40‒400) (Fig. 2F). While SIRT7(R11A, K12A) showed similar binding to nucleosomes compared to the wild-type enzyme, the N-terminal truncation in SIRT7(40‒400) reduced binding significantly (Fig. 4A,B and Supplementary Fig. 11A). Therefore, the interaction of SIRT7 with the nucleosome relies on multivalent engagement of the histone octamer beyond the acidic patch. In agreement, multiple arginine side chains along the SIRT7:octamer interface exhibit highly defined cryo-EM densities (Supplementary Figs. 5 and 10). Interestingly, this contrasts with the common arginine-anchor-based binding modes observed in other nucleosome-modifying enzymes^51^.

**Fig. 4.**
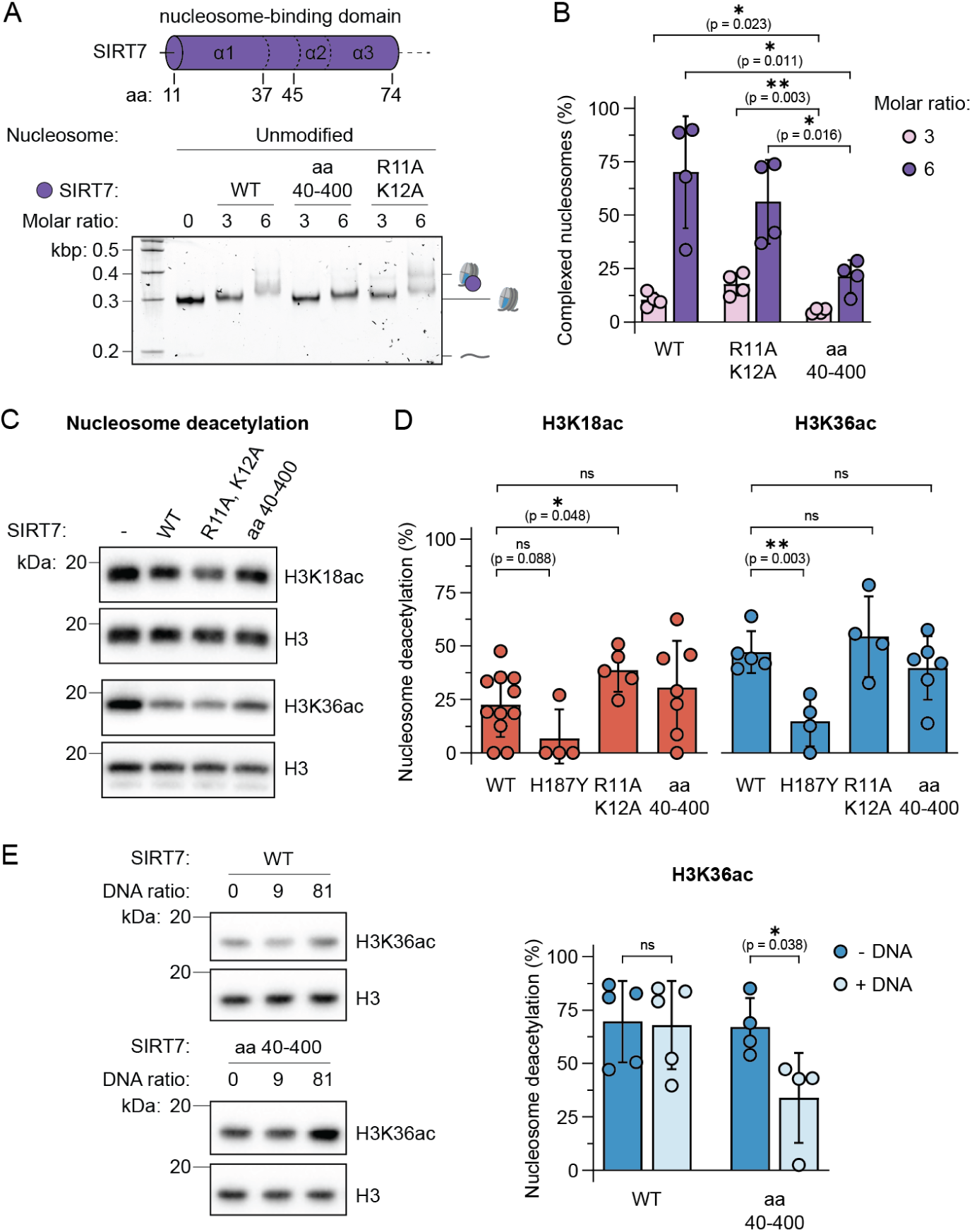
Role of the N-terminal domain to nucleosome binding and deacetylation. **A**, EMSA of unmodified nucleosomes with wild type SIRT7 and nucleosome-binding domain mutants. See Supplementary Fig. 11A for replicate gels. **B**, Quantification of SIRT7:nucleosome complex formation with unmodified nucleosomes and mutants as indicated. Error bars represent mean ± SD (*n* = 4). **C**, SIRT7 wild type and mutant activity on H3K18ac and H3K36ac nucleosomes measured by western blot. SIRT7 concentration was 50 nM and 3 nM, respectively. **E**, Quantification of nucleosome deacetylation normalized to H3 loading, at 50 nM SIRT7 for H3K18ac and 3 nM SIRT7 for H3K36ac. Error bars represent mean ± SD (*n* ≥ 4), ns: p > 0.05. **E**, Inhibitory effect of free DNA (187 bp) on SIRT7 activity, measured by western blot, and quantification of the effect of 81 equivalents of DNA, normalised to H3 loading. See Supplementary Fig. 11B for complete titration. Error bars represent mean ± SD (*n* ≥ 4), ns: p > 0.05.

We then tested the activity of these mutants, as well as an active-site mutant (H187Y)^11,57^, towards H3K18ac and H3K36ac in reconstituted nucleosomes *in vitro*. For these assays, we chose SIRT7 concentrations resulting in comparable activity towards H3K36 and H3K18 (3 and 50 nM, respectively, Fig. 4C). As expected, the active-site mutant H187Y did not exhibit much activity on either substrate (Fig. 4D)^11,57^. Conversely, the activity of SIRT7(R11A, K12A) was increased towards H3K18ac deacetylation (Fig. 4D). This finding highlights that SIRT7 does not rely on an arginine-anchor mechanism for maintaining its ability to deacetylate nucleosomes under these conditions. In contrast, mutating arginine-anchor residues in SIRT6 resulted in a loss of enzymatic activity^38,46^, further emphasising the differences in substrate recognition among the two class IV sirtuins. Interestingly, also the truncated SIRT7(40‒400) construct showed activity similar to the wild type enzyme (Fig. 4D). This can be rationalized since, under our assay conditions, dissociation from the product is the rate-limiting step for SIRT7 mononucleosome deacetylation^31^. Thus, faster substrate release of SIRT7(40‒400), due to its lower binding affinity, could be a plausible explanation for the increased deacetylation kinetics.

Based on these results, we hypothesised that the role of the SIRT7 N-terminal domain lies in its specific recruitment to nucleosomes over other nucleic acids. We thus challenged SIRT7 with increasing amounts of competitor DNA, while monitoring its H3K36 deacetylation activity on nucleosomes. Indeed, SIRT7(40‒400) was efficiently outcompeted by free DNA, while the wild-type enzyme retained its activity independent on added competitor DNA (Fig. 4E). Thus, the full nucleosome-binding domain is necessary for efficient chromatin recruitment of SIRT7 in a complex environment.

### SIRT7 site selectivity relies on specific contacts with DNA

Careful comparison of the catalytic domain conformation between the H3K36- and H3K18-bound structures revealed marked conformational changes of DNA-binding loops (Fig. 3C). To analyse the contribution of each loop to the substrate selectivity of SIRT7, we mutated key positively charged amino acids to alanine in each of them separately (loop 1: H217A, R218A; loop 2: K272A, K275A, K276A; and loop 3: R289A, R290A, Fig. 5A). Overall, the three loop-mutants showed activity on both H3K18ac and H3K36ac substrates, although with interesting differences in site selectivity (Fig. 5B).

**Fig. 5.**
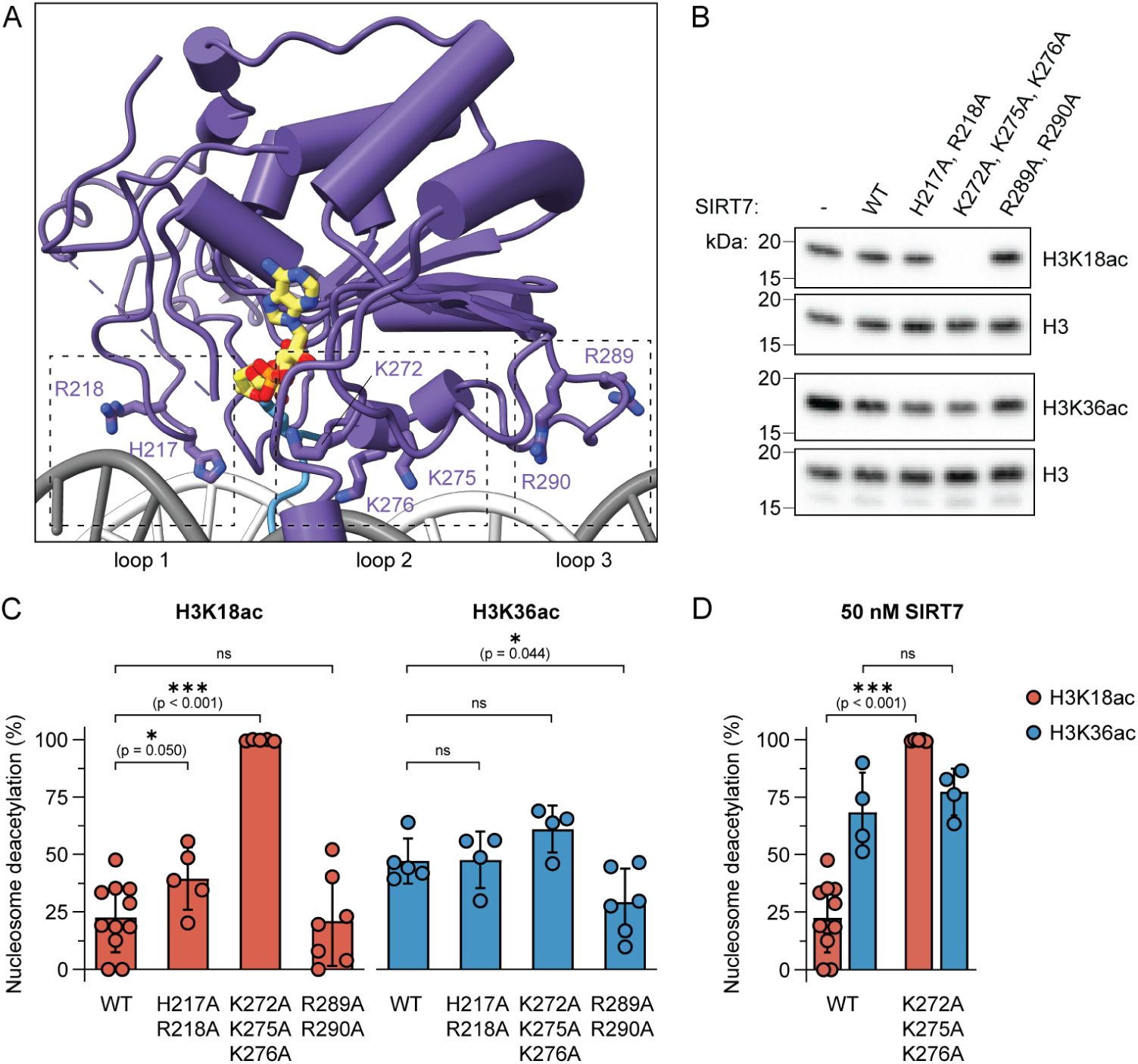
Loops flanking SIRT7 catalytic domain control substrate selectivity. **A**, Summary of mutated catalytic domain loops. **B**, SIRT7 wild type and mutant activity on H3K18ac and H3K36ac nucleosomes measured by western blot. SIRT7 concentration was 50 nM and 3 nM, respectively. **C**, Quantification of nucleosome deacetylation normalised to H3 loading, at 50 nM SIRT7 for H3K18ac and 3 nM SIRT7 for H3K36ac. See Supplementary Fig 11D for data of helix α3 mutant K72A, R73A, R74A. **D**, Quantification of wild type and K272A, K275A, K276A mutant activity at 50 nM SIRT7 concentration. Error bars represent mean ± SD (*n* ≥ 4), ns: p > 0.05.

When replacing H217 and R218 with alanine within loop 1, SIRT7 activity towards H3K18ac was increased (Fig. 5C). This indicates that binding the 2^nd^ DNA gyre on the nucleosome, thereby positioning SIRT7 over the H3 tail exit site, is suboptimal for K18 deacetylation. Releasing the enzyme from these constraints may generate a more dynamic complex with relaxed specificity.

Strikingly, mutating lysines K272, K275 and K276 in loop 2 to alanine resulted in even higher SIRT7 deacetylation activity on H3K18ac, leading to the inversion of substrate preference towards H3K18 over H3K36 (Fig. 5D). This mutant thus reveals that the loop between K272 and K276 is key for the substrate specificity of SIRT7. Based on our structures, we argue that binding of this loop to DNA forces the Kac substrate pocket to face the H3 tail exit site, where H3K36 is located, and thus makes it difficult for it to target other substrates in the H3 tail. To further test this hypothesis, we measured SIRT7 activity on H3K18ac nucleosomes lacking linker DNA (beyond SHL +7.0) and found an increase in wild type activity in line with reduced specificity (Supplementary Fig. 11C).

Finally, SIRT7 containing point mutants R289A, R290A within loop 3 showed lower activity, in particular on H3K36ac nucleosomes (Fig. 5D). Thus, linker DNA interactions are important for H3K36 targeting and contribute to the substrate selectivity.

In summary, the extended N-terminal domain of SIRT7 is crucial for chromatin recruitment, positioning the catalytic domain to tightly interact with both DNA gyres. The DNA contacts of the catalytic domain have evolved towards optimal H3K36 deacetylation and confer selectivity against other substrates. Conversely, the relaxation of DNA interactions increases SIRT7 dynamics and allows the enzyme to target H3K18. Loosening of the complex structure via mutations targeting the SIRT7:nucleosome interface further promote H3K18 deacylation activity. Together, the DNA targeting loops, in particular residues K272, 275 and K276, ensure H3K36ac selectivity, highlighting a new chromatin targeting mechanism.

## Discussion

SIRT7 is a driver of tumor growth and metastasis in various cancers and has been proposed as therapeutic target^11,14,24^. However, its complicated biochemistry, in particular the dependence on nucleosomal substrates or activation by nucleic acids, as well as the lack of structural information, has impeded the study and inhibitor development of this histone modifier.

Here, we have determined the cryo-EM structures of SIRT7 in complex with the nucleosome, targeting two key substrate residues, H3K36 and H3K18. As in other recent studies^38,48^, the use of mechanism-based inhibitor groups was crucial to stabilise the active conformation of SIRT7 contacting different target residues^33^, allowing us to identify the mechanisms of substrate selectivity in an unprecedented manner. As mechanism-based crosslinking stabilises substrate complexes regardless of their relative conversion rates^58,59^, we envision that this strategy will be widely applicable to a variety of enzyme-substrate complexes and will greatly advance the study of their mechanism of action.

Both members of class IV sirtuins, SIRT6 and SIRT7, bind nucleosomes with nanomolar affinity and are activated by these interactions^31,60^. In the case of SIRT6, the mechanism of activation is proposed to be through multivalent interaction of its disordered C-terminal domain with DNA^61^, while the catalytic domain binds the acidic patch with a SIRT6-specific arginine anchor crucial for its activity^38,46^. In contrast, here we show that SIRT7 does not rely on a single anchor, but that it establishes a multivalent interaction surface across the nucleosome via the SIRT7-specific nucleosome-binding domain. This domain is especially important for recruitment within the nucleus, as lack of contacts with the histone octamer makes SIRT7 sensitive to DNA inhibition.

While the nucleosome-binding domain forms interactions across the face of the nucleosome, the catalytic domain contacts both DNA gyres on the side of the nucleosome. This is unprecedented in chromatin-associated enzymes and key for SIRT7 selectivity towards H3K36ac substrates. Since most of the DNA contacts are made by flexible loops within the Zn^2+^-binding part of the catalytic domain^37^ and, potentially, by an unstructured region within the Rossman fold (as predicted by AlphaFold 3), we hypothesize that SIRT7 is activated by an overall conformational rearrangement of the catalytic domain when bound to the nucleosome.

While this study focuses on SIRT7 activity on nucleosomes, earlier studies have indicated that its activity can be stimulated by free nucleic acids, in particular by 5S RNA through multiple interactions with the N-terminal domain^30^. Moreover, the disordered C-terminal domain of SIRT7 has also been predicted to bind DNA and promote activation^29^. Key residues previously identified for enzyme activation by free RNA have however limited effect on nucleosome deacylation^30^. Thus, nucleosome- and RNA-dependent activation mechanisms seem to be distinct for SIRT7.

Our structures stabilised on both H3K18- and H3K36-targeting conformations enabled the detailed study of SIRT7 substrate recognition. To contact H3K36, previous structures revealed significant DNA unwrapping from the nucleosome surface^47,48^. In contrast, SIRT7 binds nucleosomes tightly just above the H3-tail exit site, with the H3 tail directly entering the active site. At the same time, SIRT7 maintains extensive contacts across both nucleosomal DNA gyres and keeps the emerging linker DNA in place, forming a ‘closed’ conformation. In contrast, when targeting H3K18, SIRT7 has to partially disengage from the nucleosome to a more ‘open’ conformation, to allow the additional amino acids 19-35 of the H3 tail loop under the bound enzyme and access the active site. In this configuration, SIRT7:DNA contacts are relaxed and the exiting linker DNA is bent, possibly due to increased space requirement of the H3 tail and interactions with a reoriented catalytic domain. Together with, and possible due to this increase in structural heterogeneity, catalytic activity drops for H3K18 deacylation.

SIRT7 H3K36 targeting thus relies on DNA contacts via multiple loops within the catalytic domain, highlighting that multivalent binding of SIRT7 to nucleosomes is optimised for H3K36 deacetylation versus other substrates. Instead of the traditional sequence-based enzyme selectivity filter, it appears that SIRT7 is selective based on its locked nucleosome-bound conformation and the distance to potential substrates. Moreover, we hypothesise that mutations in the DNA-contacting loops shift the conformational equilibrium from a ‘closed’ to a more ‘open’ state, which in turn favours enzymatic activity towards H3K18. This is indeed what we find when reengineering the enzyme. Importantly, we identified a key loop between K272 and K276 that determines SIRT7 substrate specificity, which will be important for future biochemical studies and tool development. Other reports have found that SIRT7 targets succinyl- and glutaryl-lysine PTMs within the nucleosome core (H3K122suc and H4K91glu, respectively^25,26^), which would require conformations different from those presented here. Thus, future research would determine whether the same interactions are involved in nucleosome core targeting and the required changes in chromatin structure.

The SIRT7 isozyme is present in all eukaryotes, but it only features a long helical N-terminal domain in insects and in vertebrates except for reptiles^35^. For example, in *D. melanogaster*, both N- and C-terminal tails are much longer than in humans but with a shorter N-terminal helical portion, and the “His-Arg” motif that interacts with the 2^nd^ DNA gyre is absent^35^. Based on our data, this may indicate that SIRT7 has distinct PTM selectivity in different species. Further studies will thus be necessary to investigate the role of H3K36 deacetylation throughout evolution and across species.

In conclusion, we present the structural basis of SIRT7 nucleosome deacetylation at its preferred biological substrates H3K18ac and H3K36ac, and identify the helical N-terminal fragment as nucleosome-binding domain and key loops within the catalytic domain as drivers of specificity towards H3K36ac. Moreover, the structures highlight the positioning and orientation of the active site as the drivers of substrate selectivity within the histone tails, contrary to previous models based on peptide sequence. These features are unprecedented among chromatin-binding proteins and represent a new paradigm in specific PTM targeting. The strategy used here, based on chemical modification of proteins using mechanism-based inhibitors^33^, was essential to this work. We anticipate that such approaches will further advance the study of enzyme target selection and regulation, unveiling the mechanisms behind their functional diversity.

## Methods

Complete chemical synthesis and histone purification methods can be found in the Supporting Information.

Fast protein liquid chromatography (FPLC) was performed on a ÄKTA Pure purification system equipped with UV detector and with columns as follow: HisTrap HP (5 mL, Cytiva) or HiFliQ (5 mL, ProteinArk) for Ni-NTA immobilized metal affinity chromatography (IMAC), HiTrap SP HP (5 mL, Cytiva) for ion exchange chromatography (IEX), Superdex S200 Increase 5/150 GL for analytical size-exclusion chromatography (SEC, Cytiva), Superdex S200 10/300 GL (Cytiva) for octamer SEC, and Superose 6 Increase 10/300GL (Cytiva) for SIRT7:nucleosome SEC. DNA and protein concentration was determined by UV-Vis spectroscopy, using absorption at 260 nm or 280 nm, respectively, as measured on a NanoDrop 2000 (ThermoFisher) by sample drop or a Lambda 365+ (PerkinElmer) instrument with quartz cuvettes. Protein high-resolution MS analysis was performed on a Waters Xevo G2-XS instrument equipped with UV diode array and quadrupole-time-of-flight analysis systems. Histone octamers, nucleosomes and SIRT7 mutants were analyzed on this system by HPLC-MS with a Waters Acquity BEH C4 column (1.7 µm, 100×2.10 mm, 300 Å) and gradients of eluent V (0.1% HCOOH in H2O) and eluent VI (0.1% HCOOH in MeCN) at a flow rate of 1 mL/min. Gels and western blot membranes were imaged using a ChemiDoc MP imaging system (Bio-Rad).

### Expression and purification of SIRT7 constructs

SIRT7 expression and purification was optimized from the protocol reported by Bolding *et al.*, 2023^28^. *E. coli* BL21 (DE3) cells were transformed with a pET100_D_TOPO plasmid encoding for SIRT7 or mutants thereof (generated by site-directed mutagenesis), with an N-terminal fusion to a 6 x histidine (6H) tag and a linker including a tobacco etch virus (TEV) protease cleavage site. Cells were cultured in 2-6 L autoinduction terrific broth (Formedium), or in lysogeny broth (Miller’s LB broth, Condalab) supplemented for autoinduction (6.8 g/L KH2PO4, 7.1 g/L Na2HPO4, 3.3 g/L (NH4)2SO4, 5 g/L glycerol, 0.5 g/L dextrose, 2 g/L α-lactose), and with 100 µg/mL carbenicillin or ampicillin, at 37 °C with shaking at 200 rpm until OD600 = 0.6-0.8. Then, flasks were transferred to 18 °C and shaken at 150 rpm overnight. Cell pellets were obtained by centrifugation (4000 g, 20 min, 4 °C), suspended in SIRT7 lysis buffer (20 mM Tris-HCl, 50 mM Na2HPO4, 700 mM NaCl, 1 mM EDTA, 2 mM DTT, 10% v/v glycerol, pH 7.5) with protease inhibitors (cOmplete EDTA free, Merck) and SerM nuclease A periplasmic fraction (only used for wild-type SIRT7), and flash-frozen for storage at -70 °C. Cells were lysed by sonication (10 s on and 30 s off, 60 s total time on) or by French press (3 × 15000 bar, only done for wild-type SIRT7) and centrifuged (20000 g, 20-30 min, 4 °C). The supernatants were supplemented with imidazole to a concentration of 25 mM and purified by Ni-NTA FPLC using 5→70% gradients of buffer A (20 mM Tris-HCl, 50 mM Na2HPO4, 700 mM NaCl, 1 mM EDTA, pH 7.5) and buffer B (20 mM Tris-HCl, 50 mM Na2HPO4, 700 mM NaCl, 1 mM EDTA, 500 mM imidazole, pH 7.5). Fractions containing 6xH-SIRT7 were identified by SDS-PAGE (12% acrylamide gel) and combined, supplemented with DTT (2 mM) and TEV protease (cat. # Z03030, GenScript), and dialyzed against buffer C (50 mM K2HPO4, 50 mM NaCl, 1 mM DTT, pH 7.5) at 4 °C overnight. The incubation could be done without dialysis, in which case the resulting mixture was diluted 3-5 fold with buffer C before IEX FPLC. The mixture was then purified by IEX FPLC using 0→70% gradients of buffer C and buffer D (50 mM K2HPO4, 1000 mM NaCl, 1 mM DTT, pH 7.5), and fractions containing SIRT7 were pooled, concentrated using 4 mL and 0.5 mL 30 kDa MW cut-off centrifugal filters (Amicon Ultra, Merck Millipore) to concentrations >1 µM. The solutions were supplemented with glycerol (ThermoFisher, cat. # 327255000) to 10% wt/wt concentration, aliquoted, and flash-frozen for long-term storage at -70 °C. The final samples were analyzed by UV-Vis spectroscopy and SDS-PAGE to verify purity (Supplementary Fig. 15), and by HPLC-MS to verify mutations (Supplementary Fig. 20‒21).

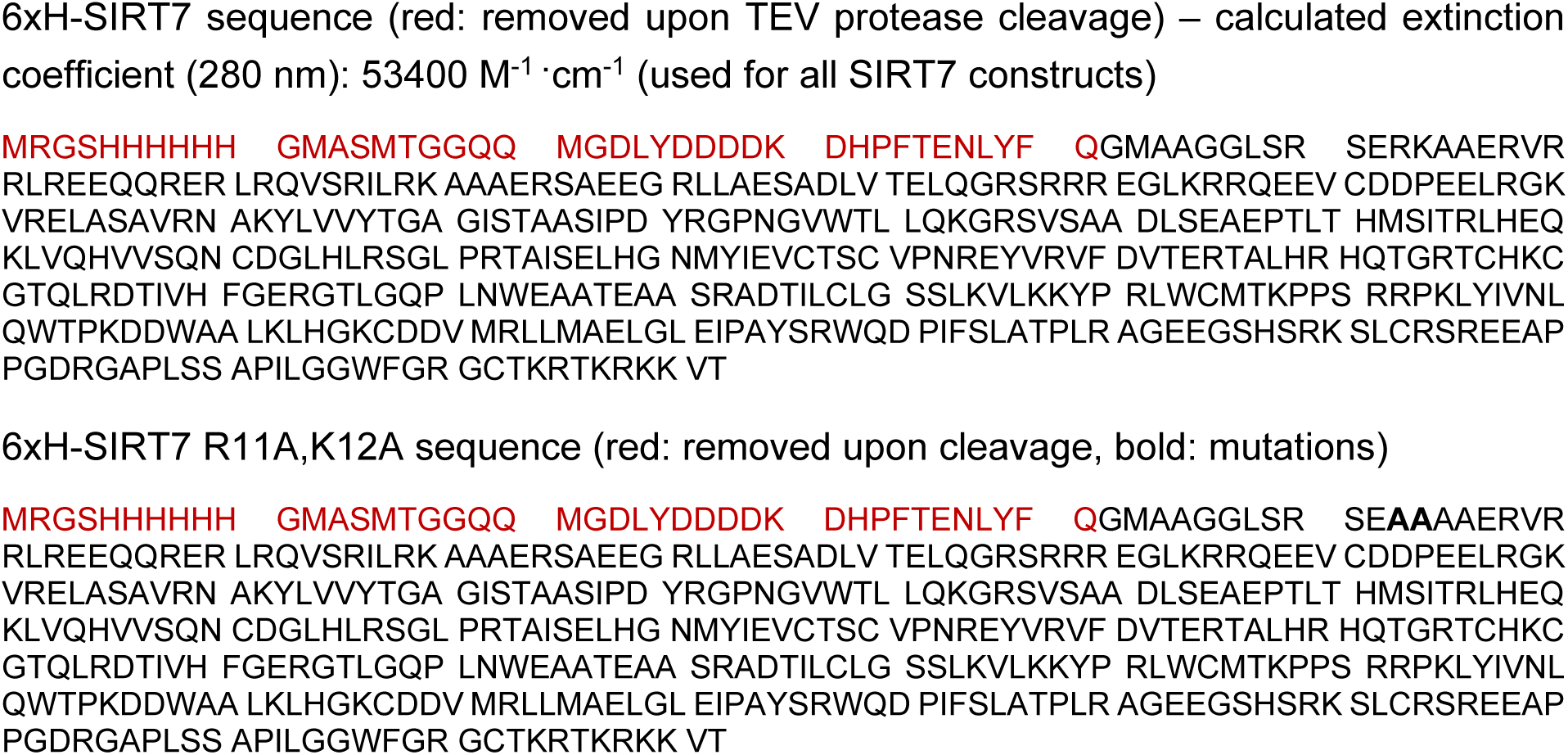
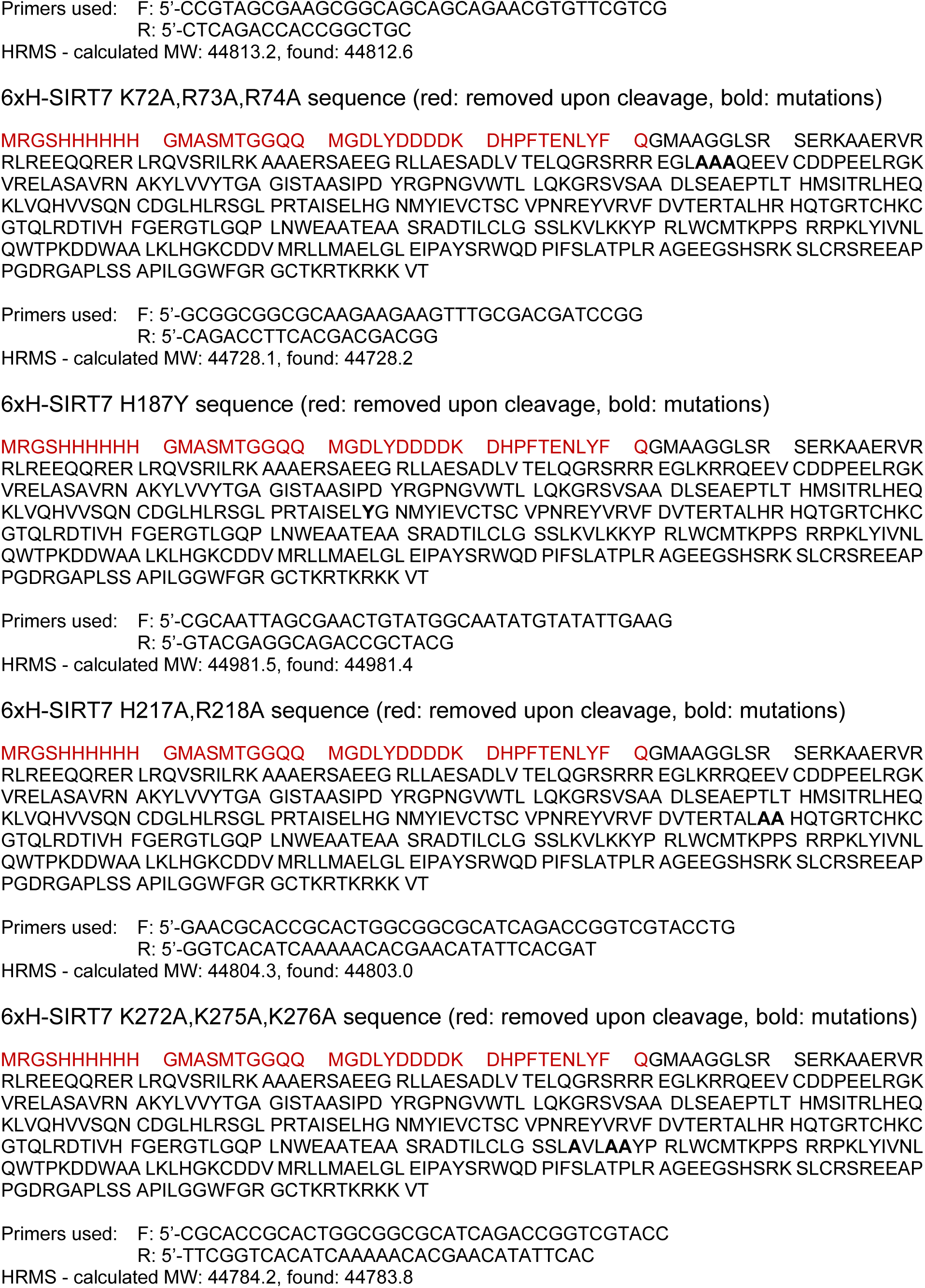
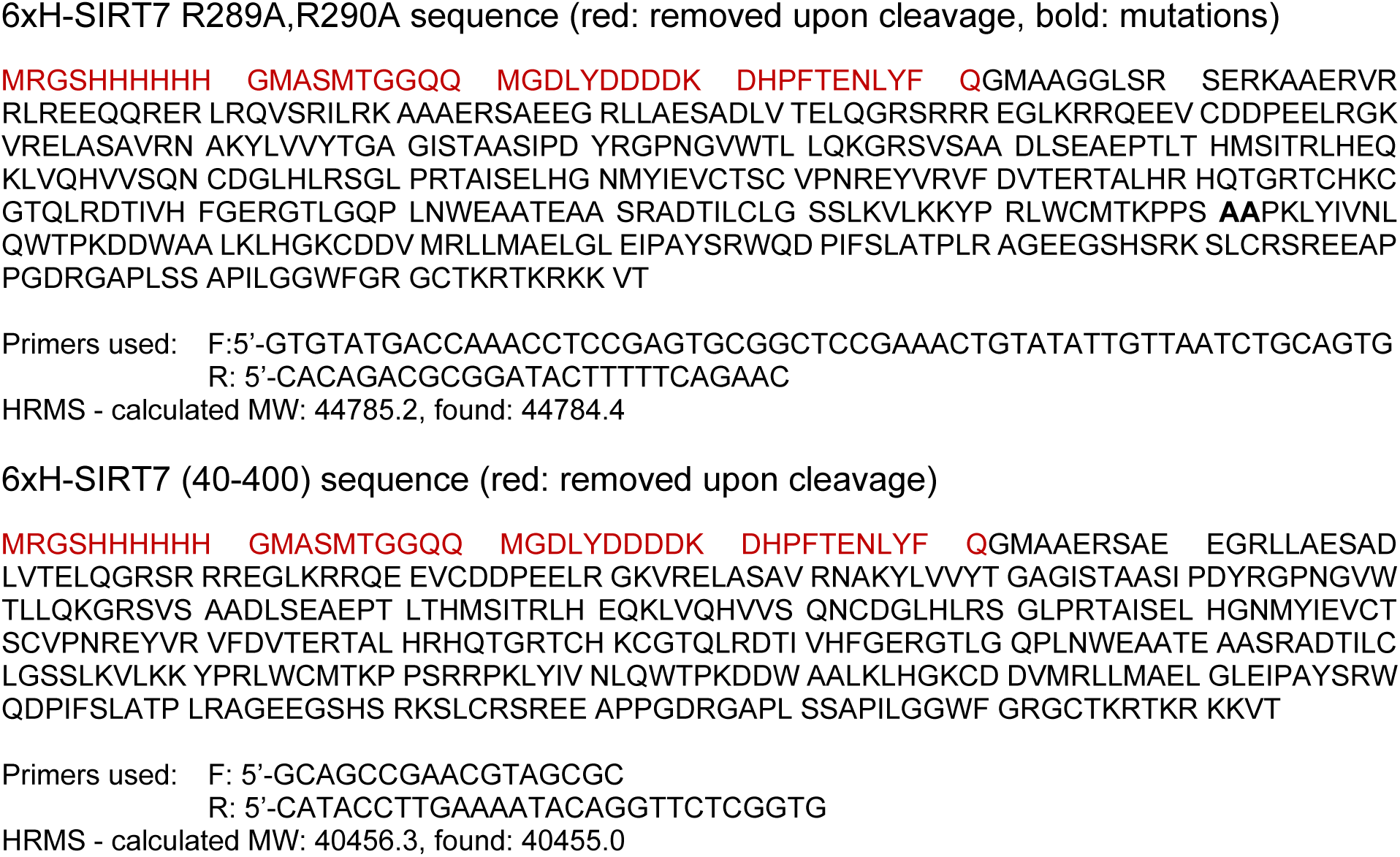

### Preparation of histone octamers

Histone octamers were prepared based on literature procedures^62^. Lyophilized histones H2A, H2B, H3 and H4 were dissolved in an unfolding buffer (6 M guanidinium chloride, 20 mM Tris-HCl, 5 mM DTT, pH 7.5) at an estimated concentration of 2 mg/mL, by vortexing and incubating for 10 min on ice. The solutions were centrifuged (21130 g, 10 min, 4 °C), and the protein concentration of the supernatants was determined by UV-Vis spectroscopy with baseline subtraction at 320 nm. The four histones were then mixed in molar ratio 1.1:1.1:1.0:1.0, and the mixture was adjusted to 1 mg/mL total protein concentration. This mixture was dialyzed in a 3.5 kDa MW cut-off membrane (Spectra/Por 3, Spectrum) three times against 3 × 0.5 L octamer buffer (10 mM Tris-HCl, 2 M NaCl, 1 mM EDTA, 1 mM DTT, pH 7.5) for 3 h, overnight, and 3 h. Thereafter, the mixture was centrifuged (21130 g, 10 min, 4 °C) and the supernatant was analyzed by UV-Vis spectroscopy and analytical HPLC (linear gradient of 0-70% B during 30 min, column #3). The supernatant was then concentrated with a 0.5 mL 30 kDa MW cut-off centrifugal filter (Amicon UItra, Merck Millipore) and purified by SEC using octamer buffer. SEC fractions were analyzed by SDS-PAGE on 15% or 17% acrylamide gels, and fractions containing equal amounts of all four histones and eluting at the expected elution volume of octamers were mixed, centrifuged (21130 g, 10 min, 4 °C), concentrated using a 0.5 mL 50 kDa MW cut-off centrifugal filter (Amicon Ultra, Merck Millipore) to concentrations >20 µM, and supplemented with glycerol (ThermoFisher, cat. # 327255000) to 50% wt/wt concentration for long-term storage at -20 °C. The final samples were analyzed by analytical HPLC and SDS-PAGE to verify equal histone amounts (Supplementary Fig. 16), and by HPLC-MS to verify histone identity and molecular integrity.

### Preparation of double-stranded DNA

Double-stranded DNA with a Widom 601 nucleosome-positioning sequence^43^ was prepared by polymerase chain reaction (PCR). Template (1.2 µg), primers (1.2 nmol each) and nucleotide mix (dNTP mix, 480 nmol each, New England Biolabs, cat. # N0447) were diluted on ice in ThermoPol reaction buffer (New England Biolabs, cat. # B9004S), and Taq DNA polymerase (120 units, New England Biolabs, cat. # M0273) was added for a total volume of 2.4 mL. The mixture was transferred to PCR tubes (50 µL/tube, Starlab, cat. # A1402-3700), and the PCR reaction was performed over 20 cycles with 1 min steps at 94 °C, 53 °C and 72 °C, and final 5 min incubation at 72 °C. The mixtures were then pooled, analyzed by agarose gel electrophoresis (Supplementary Fig. 16), diluted with buffer PB (QIAgen, cat. # 19066) and sodium acetate (0.1 M final concentration), and purified by QIAquick PCR purification (QIAgen protocol) using QIAprep 2.0 spin columns (QIAgen, cat. # 27115) onto buffer EB (QIAgen, cat. # 19086). Final samples containing 1.8-6.1 µM DNA concentration were stored at -20 °C.

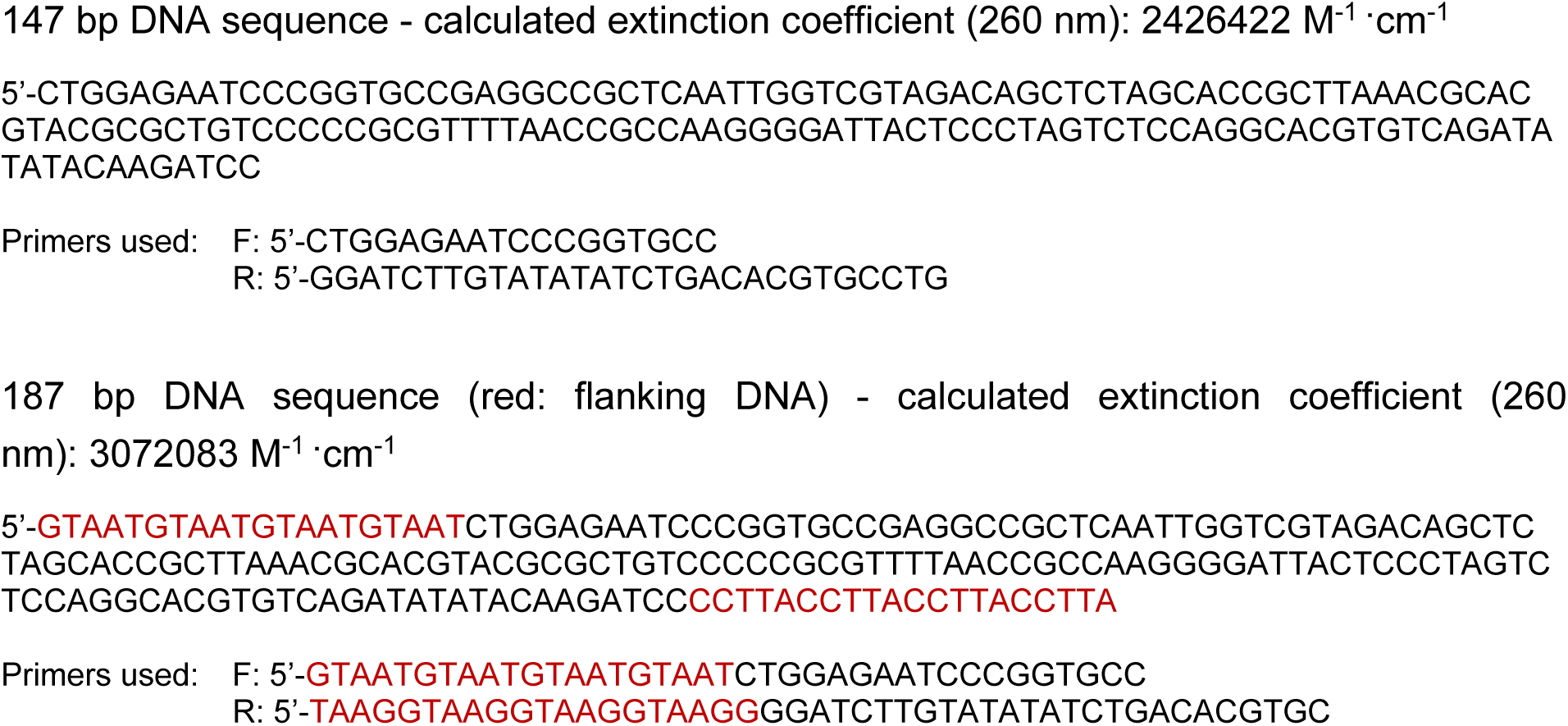

### Preparation of nucleosomes

Octamers were diluted 1:1 or 1:4 in high salt buffer (50 mM Tris-HCl, 2 M KCl, 1 mM DTT, pH 7.5) and kept on ice for 30 min. Then, double-stranded DNA (0.01-0.06 nmol, 1.0 equiv.) and diluted octamers (1.0-2.4 equiv.) were mixed in 2 M NaCl to a final volume of 30-80 µL and transferred to 10 kDa MW cut-off dialysis devices (Slide-A-Lyzer MINI, ThermoFisher Scientific, cat. # 69572). For larger preparative scales (1.0-1.2 nmol DNA, 1.4-1.5 mL final volume), samples were transferred to 3 mL dialysis cassettes (Slide-A-Lyzer, ThermoFisher Scientific, cat. # 66382). The devices were placed on 200 mL high salt buffer at 4 °C and dialyzed overnight into a low salt buffer (2 L, 50 mM Tris-HCl, 10 mM KCl, 1 mM DTT, pH 7.5) using a peristaltic pump (MiniPuls 3, Gilson). Then, samples were transferred to plastic microcentrifuge tubes, centrifuged (21130 g, 10 min, 4 °C), and analyzed by native gel electrophoresis (home-made 5% TBE gels, ran on ice at 100 V for 1.5 h with cold 0.5x TBE buffer and stained with GelRed, Supplementary Fig. 17). Titrations were performed initially to find the lowest octamer ratio at which free DNA was consumed. Then, samples prepared at such octamer ratio, and containing minimum amounts of free DNA as judged by gel electrophoresis, were combined and concentrated using 0.5 mL 50 kDa MW cut-off centrifugal filters (Amicon Ultra, Merck Millipore) to concentrations >0.8 µM (calculated based on DNA concentration). Nucleosome samples were stored at 4 °C for up to 3 months.

### Sample preparation of nucleosome-Sirt7 complexes for cryo-EM

The two complexes containing full-length SIRT7, H3K36MTU or H3K18DTU nucleosomes, and NAD^+^ were assembled by mixing SIRT7 and nucleosome at a 2.5:1 molar ratio (final concentrations: 10 μM SIRT7 and 4 μM nucleosomes) in the reconstitution buffer containing 25 mM HEPES-KOH, 50 mM KCl, and 300 μM NAD^+^, in a final volume of 300 μL. The reconstitution reaction was incubated at room temperature for 30 min and crosslinked for 10 min on ice, by adding an equal volume of the crosslinking buffer (20 mM HEPES-KOH pH 7.5, 50 mM KCl, 1 mM DTT, and 0.1% glutaraldehyde) resulting in a final glutaraldehyde concentration of 0.05%. Crosslinking was quenched by adding 60 μL of 1 M Tris pH 7.5.

For the SIRT7:H3K36MTU-nucleosome complex, 300 μL of the crosslinked complex was purified by SEC pre-equilibrated in gel filtration buffer containing 25 mM HEPES-KOH, 50 mM KCl and 1 mM DTT. The peak fractions were analysed by native gel electrophoresis on 6% Novex^TM^ TBE gels (Thermo Fisher Scientific), and fractions containing SIRT7-bound nucleosomes were pooled and concentrated to an absorbance of 3 at 280 nm for cryo-EM grid preparation (Supplementary Fig. 2A). For the SIRT7:H3K18DTU-nucleosome complex, the sample was processed by gradient fixation (GraFix)^53^ to stabilise the complex after crosslinking, instead of size exclusion chromatography. The gradient contained buffer A (10% glycerol, 25 mM HEPES-KOH, and 50 mM KCl) at the top and buffer B (40% glycerol, 25 mM HEPES-KOH, 50 mM KCl and 0.15% glutaraldehyde) at the bottom. 300 uL of the crosslinked sample was applied to the top of the gradient, followed by centrifugation (30000 rpm, 20 h, 4 °C) on a Beckman Coulter Optima XE-90 Ultracentrifuge. Subsequently, the gradient was fractionated, and crosslinking was quenched by adding a final concentration of 100 mM Tris pH 7.5. The fractions were analysed by native gel electrophoresis on 6% Novex^TM^ TBE gels, and fractions containing SIRT7-bound nucleosomes were pooled, buffer-exchanged, and concentrated to an absorbance of 3 at 280 nm for cryo-EM grid preparation (Supplementary Fig. 2B).

Cryo-EM grids for the SIRT7-nucleosome complexes were prepared by applying 3 μL of concentrated sample onto 400-gold mesh R1.2/1.3 Quantifoil grids (Quantifoil Micro Tools GmbH). These grids were rendered hydrophilic by glow discharging at 15 mA for 90 s with a PELCO easiGlow device (Ted Pella Inc.). The sample was adsorbed for 30 s on the grids at 10 °C and 100% humidity, followed by blotting and plunge-freezing into liquid ethane using a Vitrobot Mark IV plunge freezer (Thermo Fisher Scientific, TFS). Cryo-EM data were collected using the automated data acquisition software EPU (TFS) on a Titan Krios G4 transmission electron microscope (TFS), operating at 300 kV and equipped with a cold field-emission gun electron source and a Falcon4 direct detection camera. Datasets were recorded in counting mode at a physical pixel size of 0.726 Å and 0.83 Å at the sample level. Datasets were collected at a defocus range of 0.8 to 2.5 μm with a total electron dose of 50 e^−^/Å^2^. Image data were saved as Electron Event Recordings.

### Cryo-EM image processing, model building, and refinement

The cryo-EM image processing was performed using cryoSPARC v3.4^63^. The EM movie stacks were aligned and dose-weighted using patch-based motion correction (cryoSPARC implementation). Contrast transfer function (CTF) estimation was also performed using the patch-based option. For the data of the SIRT7:H3K36MTU-nucleosome complex, a blob picker was used for initial particle picking, which resulted in 792,833 particles from the initial 1000 images. These particles were used for 2D classification to generate templates for template-based particles picking on the full dataset, resulting in 7,266,736 particles. Multiple rounds of 2D classifications, ab initio reconstruction and hetero-refinement, yielded multiple 3D classes. The best 3D class comprising 1,020,323 particles was used for 3D classification with a mask covering the interaction zone between SIRT7 and the nucleosome. The best 3D classes showing distinct densities for SIRT7 were selected, resulting in 337,476 particles (Supplementary Fig. 3). Further processing of these particles by Homo refinement, Local refinement and non-uniform refinement resulted in a cryo-EM map at 2.8 Å in C1 symmetry (Supplementary Fig. 4 and Supplementary Table 1).

The 2D classes generated from the SIRT7:H3K36MTU-nucleosome complex were used for template picking for the SIRT7:H3K18DTU-nucleosome complex followed by two rounds of 2D classification, resulting in 1,053,851 particles which were subsequently used for ab-initio reconstruction. Two 3D classes containing 703,557 particles were used for hetero-refinement, followed by 3D classification with a mask covering the interaction zone between SIRT7 and the nucleosome. Further, masked 3D classification was performed on 48,563 particles obtained from the best class from the previous round of 3D classification (Supplementary Fig. 8). This resulted in a class containing complete density for the SIRT7 domain from 22,579 particles. A final map of an overall resolution of 3.4 Å in C1 symmetry was obtained following non-uniform and local refinement (Supplementary Fig. 9 and Supplementary Table 1).

Atomic models for both structures were built by docking an AlphaFold2 (ColabFold implementation) prediction of SIRT7 and the crystal structure of the human nucleosome (PDB ID: 3LZ0) into the cryo-EM map, followed by several rounds of manual rebuilding in Coot 0.9.4^64^. Real-space refinement for all built models was performed using Phenix, version 1.19.2-4158^65^, using a general restraints setup (Supplementary Table 1).

### Structure visualization and analysis

Structural alignments and superpositions were performed using UCSF Chimera, UCSF ChimeraX v1.4^66^ and PyMOL Version 1.8.2.0. Gel images were processed and prepared on ImageJ (version 1.53k)^67^. Figures were rendered using UCSF ChimeraX.

### Determination of SIRT7 concentration

SIRT7 aliquots were thawed, centrifuged (21130 g, 3 min, 4 °C), and the concentration of the supernatant was estimated by Nanodrop. Then, ∼0.25 µg protein were analyzed by SDS-PAGE (SurePAGE 4-12% gels, Genscript cat. # M00654, ran at 100 V and stained with Coomasie blue) together with a set of BSA standard dilutions (TFS, cat. # 23209), and bands were quantified using ImageJ. BSA data were analyzed to generate a standard curve with r^2^ > 0.98, which was then used to calculate SIRT7 concentration. Sample preparation and SDS-PAGE was repeated twice, and the average of calculated SIRT7 concentrations was used for all further experiments.

### Electrophoretic mobility shift assays (EMSAs)

Nucleosomes (100 nM), NAD^+^ (0 or 150 µM) and SIRT7 (0‒1.6 µM, as indicated on each experiment) were mixed in 0.5 mL low-binding microcentrifuge tubes in EMSA buffer (10 mM Tris-HCl, 50 mM KCl, 1 mM DTT, pH 7.5) for a final volume of 8 µL while kept on ice. Tubes were then transferred to a thermal shaker at 25 °C and incubated for 30 min at 800 rpm, followed by cooling on ice, addition of 50% wt/v glycerol in EMSA buffer (2 µL, 10% final glycerol concentration), and analysis by native gel electrophoresis (home-made 5% TBE gels, ran on ice at 100 V for 2 h with cold 0.25x TBE buffer and stained with GelRed). Bands were quantified from raw TIF files using ImageJ, and data was analyzed using GraphPad Prism 10. Groups were compared by unpaired t-test for statistical analysis.

### Deacetylation assays with Western blot readout

Nucleosomes (200 nM), NAD^+^ (500 µM) and SIRT7 (0‒0.8 µM) were mixed in 0.5 mL low-binding microcentrifuge tubes in deacylation buffer (10 mM Tris-HCl, 50 mM KCl, 1 mM DTT, 0.2 mg/mL BSA, pH 7.5) for a final volume of 8 µL while kept on ice. Then, tubes were transferred to a thermal shaker at 37 °C and incubated for 2 h at 800 rpm, followed by cooling on ice, mixing with 4x Laemmli sample buffer (2.7 µL, Bio-Rad, cat. #1610747, supplemented with β-mercaptoethanol), and separation by SDS-PAGE (home-made 12% or 17% gels, ran at 180 V for 0.5‒1 h with Tris-glycine buffer). Gels were shortly incubated in 20% EtOH, and the samples were transferred to polyvinylidene difluoride membranes (PVDF, ThermoFisher Scientific, cat. # IB24001) using an iBlot2 western blot transfer system (ThermoFisher Scientific). After transfer, membranes were blocked in 5% skimmed milk in TBST for 1 h at room temperature, washed with TBST (2 × 1 min), and incubated with primary antibodies at 4 °C overnight (1:1000 or 1:2000 dilution in TBST with 2% BSA and 0.02% NaN3). Then, membranes were washed with TBST (3 × 1 min), incubated with horseradish peroxidase (HRP)-conjugated secondary antibodies for 1 h at room temperature (1:10000 dilution in TBST with 2% BSA), and washed with TBST (2 × 1 min) and TBS (1 × 1 min) before imaging using Clarity western enhanced chemiluminescence substrate (Bio-Rad, cat. # 1705061). Membranes were always stained first with H3 (C-term) before any Kac staining. Bands were quantified from TIF raw files using ImageJ, and data was normalized to the H3 signal and analyzed using GraphPad Prism 10. Groups were compared by unpaired t-test for statistical analysis.

The following antibodies were used:

| Target | Organism | Vendor | Cat. # | Dilution |
| --- | --- | --- | --- | --- |
| H3K18ac | Rabbit | ThermoFisher Scientific | 703896 | 1:1000 |
| H3K36ac | Rabbit | Sigma-Aldrich | SAB5600229 | 1:1000 |
| H3 (C-term) | Mouse | Cell Signaling Technology | 3638 | 1:2000 |
| SIRT7 (N-term) | Rabbit | Cell Signaling Technology | 5360 | 1:1000 |
| Mouse IgG<br>(HRP-conjugated) | Horse | Cell Signaling Technology | 7076 | 1:10000 |
| Rabbit IgG<br>(HRP-conjugated) | Goat | Cell Signaling Technology | 7074 | 1:10000 |

### Deacylation assays with mass spectrometry readout

Nucleosomes (200 nM), NAD^+^ (500 µM) and SIRT7 (0‒0.2 µM) were mixed in 0.5 mL low-binding microcentrifuge tubes in deacylation buffer (10 mM Tris-HCl, 50 mM KCl, 1 mM DTT, 0.2 mg/mL BSA, pH 7.5) for a final volume of 8 µL while kept on ice. Then, tubes were transferred to a thermal shaker at 37 °C and incubated for 2 h at 800 rpm, followed by cooling on ice and mixing with 3.3% formic acid in H2O (12 µL, 2% final concentration). Samples were mixed by vortexing, centrifuged (21130 g, 3 min, 4 °C), and analysed by HPLC-MS on a Waters Xevo G2-XS instrument. Samples were run through a Waters Acquity BEH C4 column (1.7 µm, 100×2.10 mm, 300 Å) using 15 min gradients of eluent V and eluent VI at a flow rate of 1 mL/min. Then, chromatograms of the indicated ions were extracted and exported to GraphPad Prism 10, where they were smoothed, integrated, and analyzed. Baseline correction was performed against control nucleosomes (for acylated substrate analysis) or against reactions without SIRT7 (for deacylated product analysis).

The following ion chromatograms were extracted:

| Nucleosome | Type of analysis | Ion identity | m/z |
| --- | --- | --- | --- |
| H3K18ac | Substrate | [M + 21H]<sup>21+</sup> | 729.52 |
| H3K18ac | Product | [M + 21H]<sup>21+</sup> | 727.52 |
| H3K18dec | Substrate | $[M + 21H]^{21+}$ | 734.86 |
| H3K18dec | Product | $[M + 21H]^{21+}$ | 727.52 |
| H3K36ac | Substrate | $[M + 21H]^{21+}$ | 731.05 |
| H3K36ac | Product | $[M + 21H]^{21+}$ | 729.04 |
| H3K36dec | Substrate | $[M + 21H]^{21+}$ | 736.39 |
| H3K36dec | Product | $[M + 21H]^{21+}$ | 729.04 |

## Acknowledgements

We would like to thank Dr. Julie Bolding and Prof. Christian A. Olsen for sharing of reagents, Dr. Kelvin Lau for assistance with the optimization of SIRT7 production, and Prof. Thomas Schalch for comments on the manuscript. The cryo-EM data collection was performed at the Dubochet Center for Imaging Lausanne (a common initiative from EPFL, UNIGE, UNIL, UNIBE) with the help of A. Myasnikov, B. Beckert, S. Nazarov, I. Mohammed and E. Uchikawa. This work was supported by the Independent Research Fund Denmark (DFF International Postdoctoral Grant #2028.00011B to C.M.-Y.), the Swiss National Science Foundation (SNSF Swiss Postdoctoral Fellowship #TMPFP2_217187 to C.M.-Y., and grant IZLCZO_206089 to D.N. and H.S.), “la Caixa” Foundation (ID 100010434, fellowship LCF/BQ/EU22/11930058 to E.C.-S), and EPFL.

## Author contributions

Conceptualization, C.M.-Y. and B.F.; Methodology, C.M.-Y., B.E.E. and E.C.-S.;

Investigation, C.M.-Y., B.E.E., P.N.F. and D.N.; Resources, C.M.-Y. and E.C.-S.; Formal Analysis, C.M.-Y., B.E.E., D.N. and B.F.; Writing – Original Draft, C.M.-Y.; Writing – Review & Editing, all authors; Visualization, C.M.-Y. and B.E.E.; Supervision, C.M.-Y., H.S. and B.F.; Funding Acquisition, C.M.-Y., H.S. and B.F.

## Additional information

Supplementary information contains supplementary figures and tables, detailed methods of chemical synthesis and purification of histone constructs, and analytical data of all peptide, protein and DNA materials.

## References

1. Choudhary, C., Weinert, B.T., Nishida, Y., Verdin, E. & Mann, M. The growing landscape of lysine acetylation links metabolism and cell signalling. Nat Rev Mol Cell Biol 15, 536–550 (2014).

2. Wang, M. & Lin, H. Understanding the Function of Mammalian Sirtuins and Protein Lysine Acylation. Annu Rev Biochem 90, 245–285 (2021).

3. Jing, H. & Lin, H. Sirtuins in epigenetic regulation. Chem Rev 115, 2350–75 (2015).

4. Frye, R.A. Phylogenetic classification of prokaryotic and eukaryotic Sir2-like proteins. Biochem Biophys Res Commun 273, 793–8 (2000).

5. Gomes, P., Leal, H., Mendes, A.F., Reis, F. & Cavadas, C. Dichotomous Sirtuins: Implications for Drug Discovery in Neurodegenerative and Cardiometabolic Diseases. Trends Pharmacol Sci 40, 1021–1039 (2019).

6. Chalkiadaki, A. & Guarente, L. The multifaceted functions of sirtuins in cancer. Nat Rev Cancer 15, 608–24 (2015).

7. Ford, E. et al. Mammalian Sir2 homolog SIRT7 is an activator of RNA polymerase I transcription. Genes Dev 20, 1075–1080 (2006).

8. Grob, A. et al. Involvement of SIRT7 in resumption of rDNA transcription at the exit from mitosis. J Cell Sci 122, 489–498 (2009).

9. Ianni, A., Hoelper, S., Krueger, M., Braun, T. & Bober, E. Sirt7 stabilizes rDNA heterochromatin through recruitment of DNMT1 and Sirt1. Biochem Biophys Res Commun 492, 434–440 (2017).

10. Paredes, S. et al. The epigenetic regulator SIRT7 guards against mammalian cellular senescence induced by ribosomal DNA instability. J Biol Chem 293, 11242–11250 (2018).

11. Barber, M.F. et al. SIRT7 links H3K18 deacetylation to maintenance of oncogenic transformation. Nature 487, 114–118 (2012).

12. Malik, S. et al. SIRT7 inactivation reverses metastatic phenotypes in epithelial and mesenchymal tumors. Sci Rep 5, 9841 (2015).

13. Zhang, C. et al. CYP2E1-dependent upregulation of SIRT7 is response to alcohol mediated metastasis in hepatocellular carcinoma. Cancer Gene Ther 29, 1961–1974 (2022).

14. Chen, K.L. et al. SIRT7 depletion inhibits cell proliferation, migration, and increases drug sensitivity by activating p38MAPK in breast cancer cells. J Cell Physiol 233, 6767–6778 (2018).

15. Kumari, P. et al. SIRT7 promotes lung cancer progression by destabilizing the tumor suppressor ARF. Proc Natl Acad Sci U S A 121, e2409269121 (2024).

16. Vazquez, B.N. et al. SIRT7 mediates L1 elements transcriptional repression and their association with the nuclear lamina. Nucleic Acids Research 47, 7870–7885 (2019).

17. Wu, S. & Jia, S. Functional Diversity of SIRT7 Across Cellular Compartments: Insights and Perspectives. Cell Biochem Biophys 81, 409–419 (2023).

18. Paredes, S., Villanova, L. & Chua, K.F. Molecular pathways: emerging roles of mammalian Sirtuin SIRT7 in cancer. Clin Cancer Res 20, 1741–1746 (2014).

19. Wang, W.W. et al. A Click Chemistry Approach Reveals the Chromatin-Dependent Histone H3K36 Deacylase Nature of SIRT7. J Am Chem Soc 141, 2462–2473 (2019).

20. Wang, Z. et al. Combinatorial patterns of histone acetylations and methylations in the human genome. Nat Genet 40, 897–903 (2008).

21. Pfister, S.X. et al. SETD2-dependent histone H3K36 trimethylation is required for homologous recombination repair and genome stability. Cell Rep 7, 2006–18 (2014).

22. Tang, X. et al. SIRT7 antagonizes TGF-beta signaling and inhibits breast cancer metastasis. Nat Commun 8, 318 (2017).

23. Vazquez, B.N. et al. SIRT7 and p53 interaction in embryonic development and tumorigenesis. Front Cell Dev Biol 11, 1281730 (2023).

24. Ianni, A., Kumari, P., Tarighi, S., Braun, T. & Vaquero, A. SIRT7: a novel molecular target for personalized cancer treatment? Oncogene (2024).

25. Li, L. et al. SIRT7 is a histone desuccinylase that functionally links to chromatin compaction and genome stability. Nat Commun 7, 12235 (2016).

26. Bao, X. et al. Glutarylation of Histone H4 Lysine 91 Regulates Chromatin Dynamics. Mol Cell 76, 660–675 e9 (2019).

27. Simonet, N.G. et al. SirT7 auto-ADP-ribosylation regulates glucose starvation response through mH2A1. Sci Adv 6, eaaz2590 (2020).

28. Bolding, J.E. et al. Substrates and Cyclic Peptide Inhibitors of the Oligonucleotide-Activated Sirtuin 7. Angew Chem Int Ed Engl 62, e202314597 (2023).

29. Tong, Z. et al. SIRT7 Is Activated by DNA and Deacetylates Histone H3 in the Chromatin Context. ACS Chem Biol 11, 742–7 (2016).

30. Tong, Z. et al. SIRT7 Is an RNA-Activated Protein Lysine Deacylase. ACS Chem Biol 12, 300–310 (2017).

31. Kuznetsov, V.I., Liu, W.H., Klein, M.A. & Denu, J.M. Potent Activation of NAD(+)-Dependent Deacetylase Sirt7 by Nucleosome Binding. ACS Chem Biol 17, 2248–2261 (2022).

32. Tanabe, K. et al. LC-MS/MS-based quantitative study of the acyl group- and site-selectivity of human sirtuins to acylated nucleosomes. Sci Rep 8, 2656 (2018).

33. Moreno-Yruela, C. & Fierz, B. Revealing chromatin-specific functions of histone deacylases. Biochem Soc Trans (2024).

34. Kiran, S. et al. Intracellular distribution of human SIRT7 and mapping of the nuclear/nucleolar localization signal. FEBS J 280, 3451–3466 (2013).

35. Lagunas-Rangel, F.A. Bioinformatic analysis of SIRT7 sequence and structure. J Biomol Struct Dyn 41, 8081–8091 (2023).

36. Priyanka, A., Solanki, V., Parkesh, R. & Thakur, K.G. Crystal structure of the N-terminal domain of human SIRT7 reveals a three-helical domain architecture. Proteins 84, 1558–1563 (2016).

37. Rajabi, N., Galleano, I., Madsen, A.S. & Olsen, C.A. Targeting Sirtuins: Substrate Specificity and Inhibitor Design. Prog Mol Biol Transl Sci 154, 25–69 (2018).

38. Wang, Z.A. et al. Structural Basis of Sirtuin 6-Catalyzed Nucleosome Deacetylation. J Am Chem Soc 145, 6811–6822 (2023).

39. Rajabi, N., Nielsen, A.L. & Olsen, C.A. Dethioacylation by Sirtuins 1-3: Considerations for Drug Design Using Mechanism-Based Sirtuin Inhibition. ACS Med Chem Lett 11, 1886–1892 (2020).

40. Hirsch, B.M. et al. A mechanism-based potent sirtuin inhibitor containing Nepsilon-thiocarbamoyl-lysine (TuAcK). Bioorg Med Chem Lett 21, 4753–4757 (2011).

41. Troelsen, K.S. et al. Mitochondria-targeted inhibitors of the human SIRT3 lysine deacetylase. RSC Chem Biol 2, 627–635 (2021).

42. Guidotti, N. & Fierz, B. Semisynthesis and Reconstitution of Nucleosomes Carrying Asymmetric Histone Modifications. Methods Mol Biol 2133, 263–291 (2020).

43. Lowary, P.T. & Widom, J. New DNA sequence rules for high affinity binding to histone octamer and sequence-directed nucleosome positioning. J Mol Biol 276, 19–42 (1998).

44. Nielsen, A.L. et al. Mechanism-based inhibitors of SIRT2: structure–activity relationship, X-ray structures, target engagement, regulation of α-tubulin acetylation and inhibition of breast cancer cell migration. RSC Chem Biol 2, 612–626 (2021).

45. Zhou, Y. et al. The bicyclic intermediate structure provides insights into the desuccinylation mechanism of human sirtuin 5 (SIRT5). J Biol Chem 287, 28307–14 (2012).

46. Chio, U.S. et al. Cryo-EM structure of the human Sirtuin 6-nucleosome complex. Sci Adv 9, eadf7586 (2023).

47. Li, W. et al. Molecular basis of nucleosomal H3K36 methylation by NSD methyltransferases. Nature 590, 498–503 (2021).

48. Spangler, C.J. et al. Structural basis of paralog-specific KDM2A/B nucleosome recognition. Nat Chem Biol 19, 624–632 (2023).

49. Wang, H., Farnung, L., Dienemann, C. & Cramer, P. Structure of H3K36-methylated nucleosome-PWWP complex reveals multivalent cross-gyre binding. Nat Struct Mol Biol 27, 8–13 (2020).

50. McGinty, R.K. & Tan, S. Nucleosome structure and function. Chem Rev 115, 2255–73 (2015).

51. McGinty, R.K. & Tan, S. Principles of nucleosome recognition by chromatin factors and enzymes. Curr Opin Struct Biol 71, 16–26 (2021).

52. Abramson, J. et al. Accurate structure prediction of biomolecular interactions with AlphaFold 3. Nature (2024).

53. Stark, H. GraFix: stabilization of fragile macromolecular complexes for single particle cryo-EM. Methods Enzymol 481, 109–26 (2010).

54. Vogler, C. et al. Histone H2A C-terminus regulates chromatin dynamics, remodeling, and histone H1 binding. PLoS Genet 6, e1001234 (2010).

55. Zhou, B.R. et al. Distinct Structures and Dynamics of Chromatosomes with Different Human Linker Histone Isoforms. Mol Cell 81, 166–182 e6 (2021).

56. Arimura, Y., Shih, R.M., Froom, R. & Funabiki, H. Structural features of nucleosomes in interphase and metaphase chromosomes. Mol Cell 81, 4377–4397 e12 (2021).

57. Hoff, K.G., Avalos, J.L., Sens, K. & Wolberger, C. Insights into the sirtuin mechanism from ternary complexes containing NAD+ and acetylated peptide. Structure 14, 1231–40 (2006).

58. Wu, M. et al. Lysine-14 acetylation of histone H3 in chromatin confers resistance to the deacetylase and demethylase activities of an epigenetic silencing complex. Elife 7(2018).

59. Moreno-Yruela, C. et al. Hydroxamic acid-modified peptide microarrays for profiling isozyme-selective interactions and inhibition of histone deacetylases. Nat Commun 12, 62 (2021).

60. Gil, R., Barth, S., Kanfi, Y. & Cohen, H.Y. SIRT6 exhibits nucleosome-dependent deacetylase activity. Nucleic Acids Res 41, 8537–8545 (2013).

61. Liu, W.H. et al. Multivalent interactions drive nucleosome binding and efficient chromatin deacetylation by SIRT6. Nat Commun 11, 5244 (2020).

62. Dyer, P.N. et al. Reconstitution of nucleosome core particles from recombinant histones and DNA. Methods Enzymol 375, 23–44 (2004).

63. Punjani, A., Rubinstein, J.L., Fleet, D.J. & Brubaker, M.A. cryoSPARC: algorithms for rapid unsupervised cryo-EM structure determination. Nat Methods 14, 290–296 (2017).

64. Emsley, P., Lohkamp, B., Scott, W.G. & Cowtan, K. Features and development of Coot. Acta Crystallogr D Biol Crystallogr 66, 486–501 (2010).

65. Adams, P.D. et al. PHENIX: a comprehensive Python-based system for macromolecular structure solution. Acta Crystallogr D Biol Crystallogr 66, 213–21 (2010).

66. Meng, E.C. et al. UCSF ChimeraX: Tools for structure building and analysis. Protein Sci 32, e4792 (2023).

67. Schindelin, J., et al. Fiji: an open-source platform for biological-image analysis. Nat Methods 9, 676-82 (2012).

