## Supplementary Information for "Structural basis of SIRT7 nucleosome engagement and substrate specificity"

### Supplementary Figures 1-11

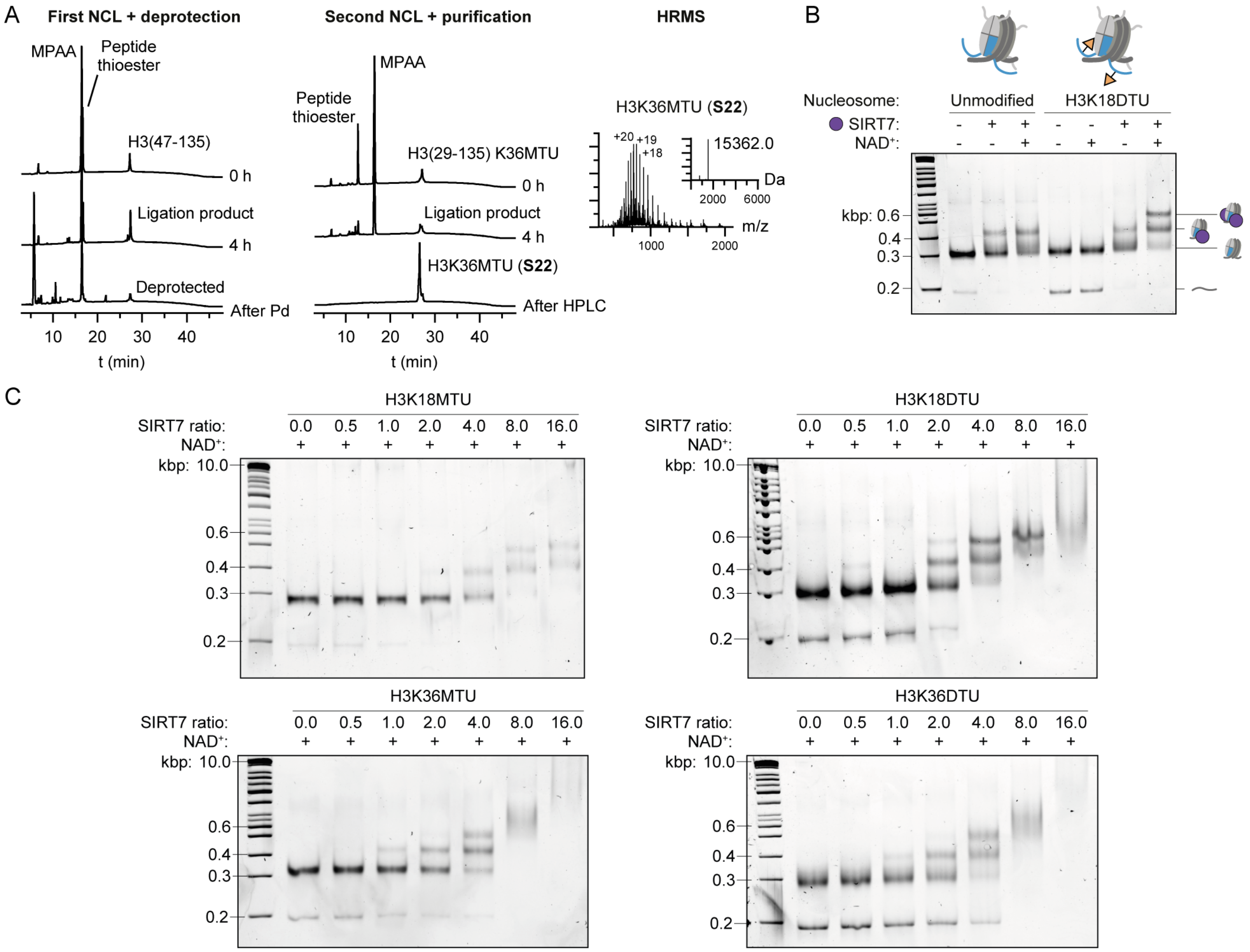

**Supplementary Fig. 1. Synthesis and testing of thiourea-functionalized nucleosomes. A**, Summary of full length H3K36MTU (**S22**) semi-synthesis. Analytical HPLC traces of the reaction mixture during the first NCL reaction and subsequent thiazolidine deprotection (left), analytical HPLC traces of the reaction mixture during the second NCL reaction and after final purification (center), and final MS spectrum and deconvoluted mass of the final purified product (right). See Supplementary Figs.18, 19, 22, 23 for complete purity traces and MS spectra of all synthesized histones. **B**, SIRT7 EMSA of nucleosome samples with and without H3K18DTU modification (SIRT7-nucleosome ratio: 3.0). **C**, Representative dose-response EMSAs with each of the four thiourea-containing nucleosomes prepared.

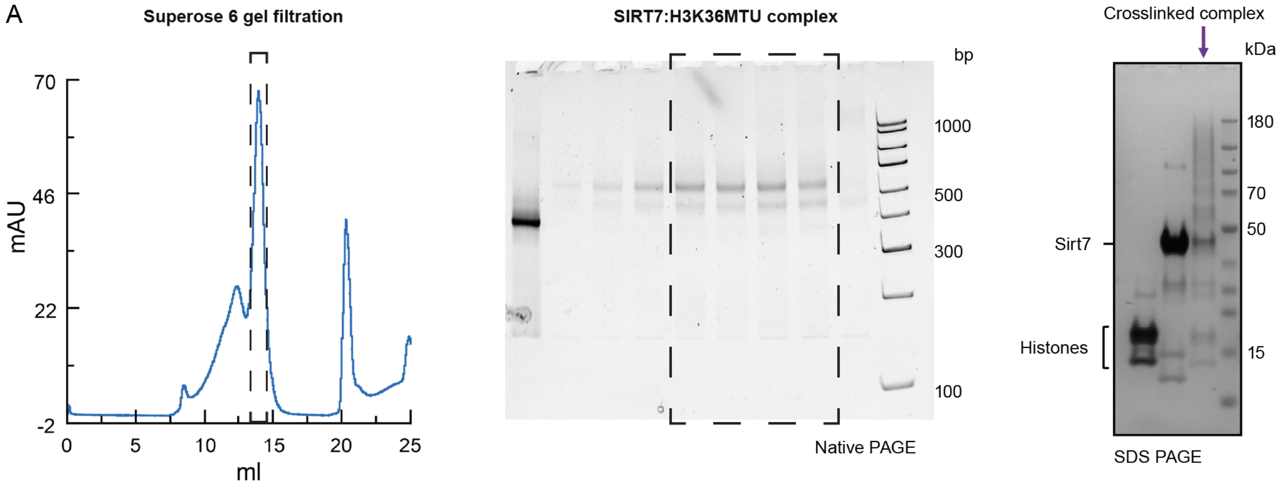

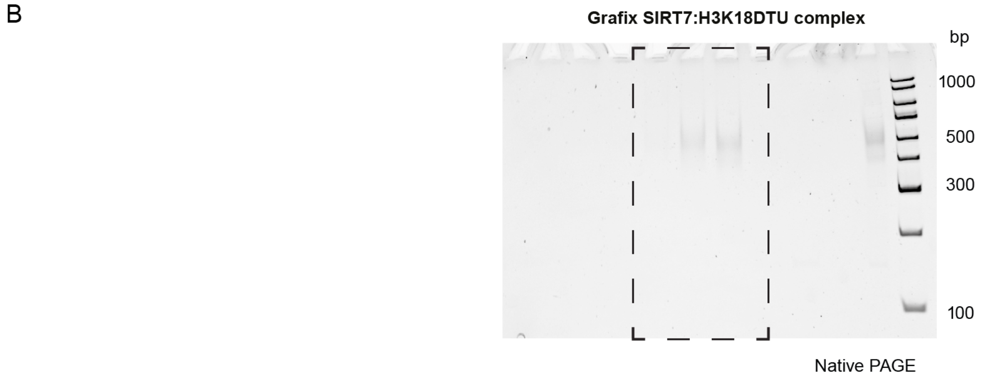

**Supplementary Fig. 2. Purification of SIRT7:nucleosome complexes. A**, Left panel shows the gel filtration chromatogram from the purification of the crosslinked SIRT7:H3K36-nucleosome complex, the middle panel shows a Native gel of the gel filtration fraction and the right panel shows an SDS-PAGE gel of the pooled crosslinked complex. Dashed lines indicate the fractions pooled for cryo-EM. **B**, Native gel of fractions from Grafix purification of the SIRT7:H3K18DTU nucleosome complex. Dashed lines indicate the fractions pooled for cryo-EM.

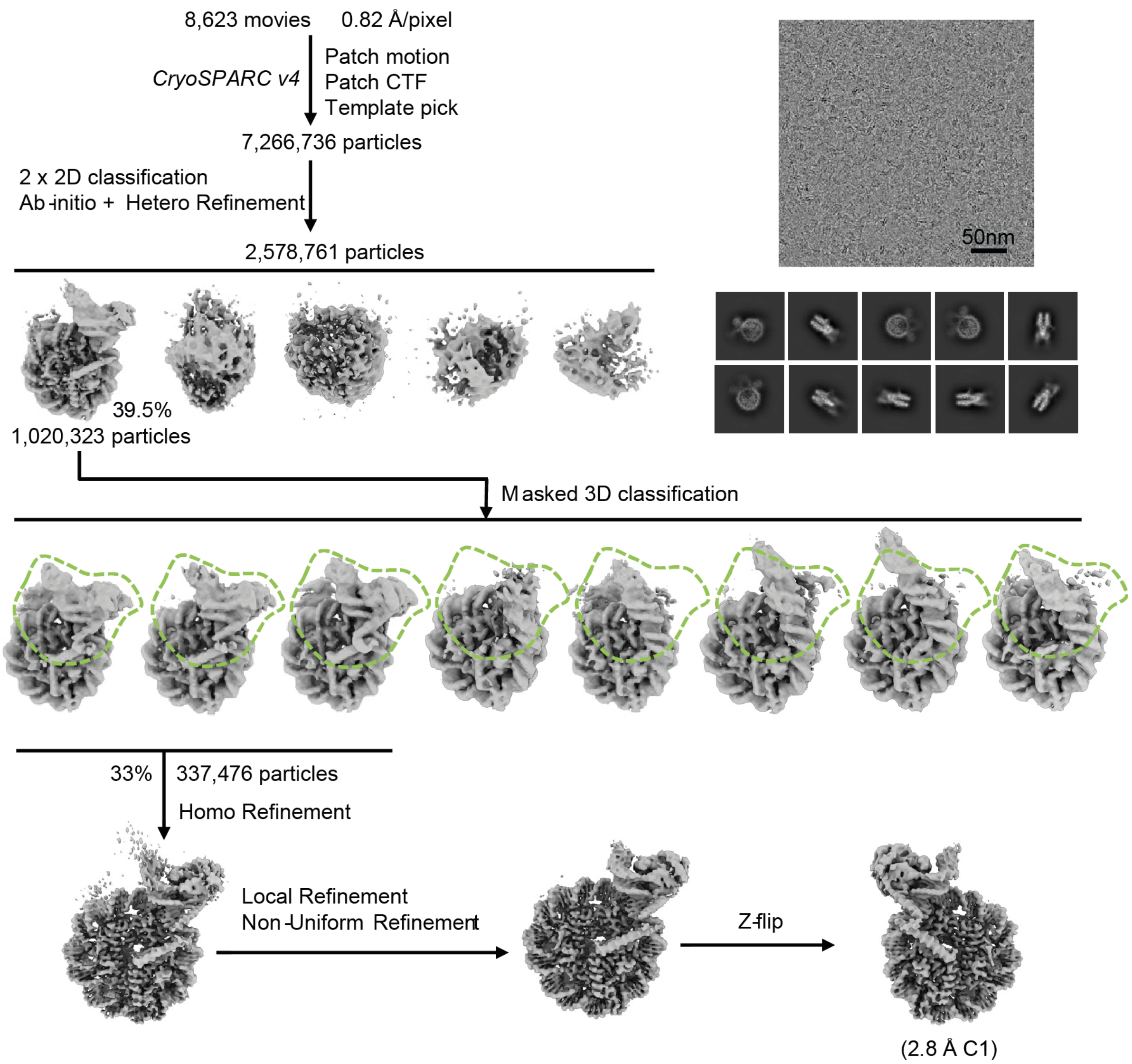

**Supplementary Fig. 3. SIRT7:H3K36MTU complex data processing workflow.** Processing workflow for SIRT7:H3K36MTU complex showing 2D class averages, 3D classification and refinement steps. Focused classification with the regions covered by the mask are also shown.

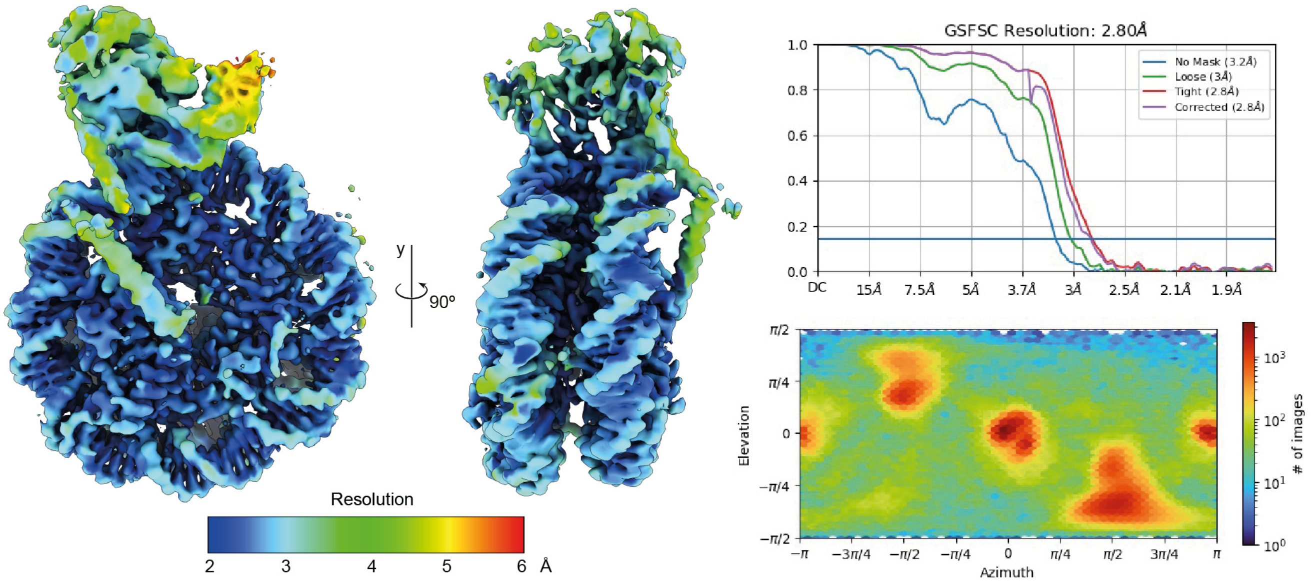

**Supplementary Fig. 4.** Local resolution map, Fourier Shell Correlation (FSC) curve and particle distribution orientation of the SIRT7:H3K36MTU-nucleosome complex.

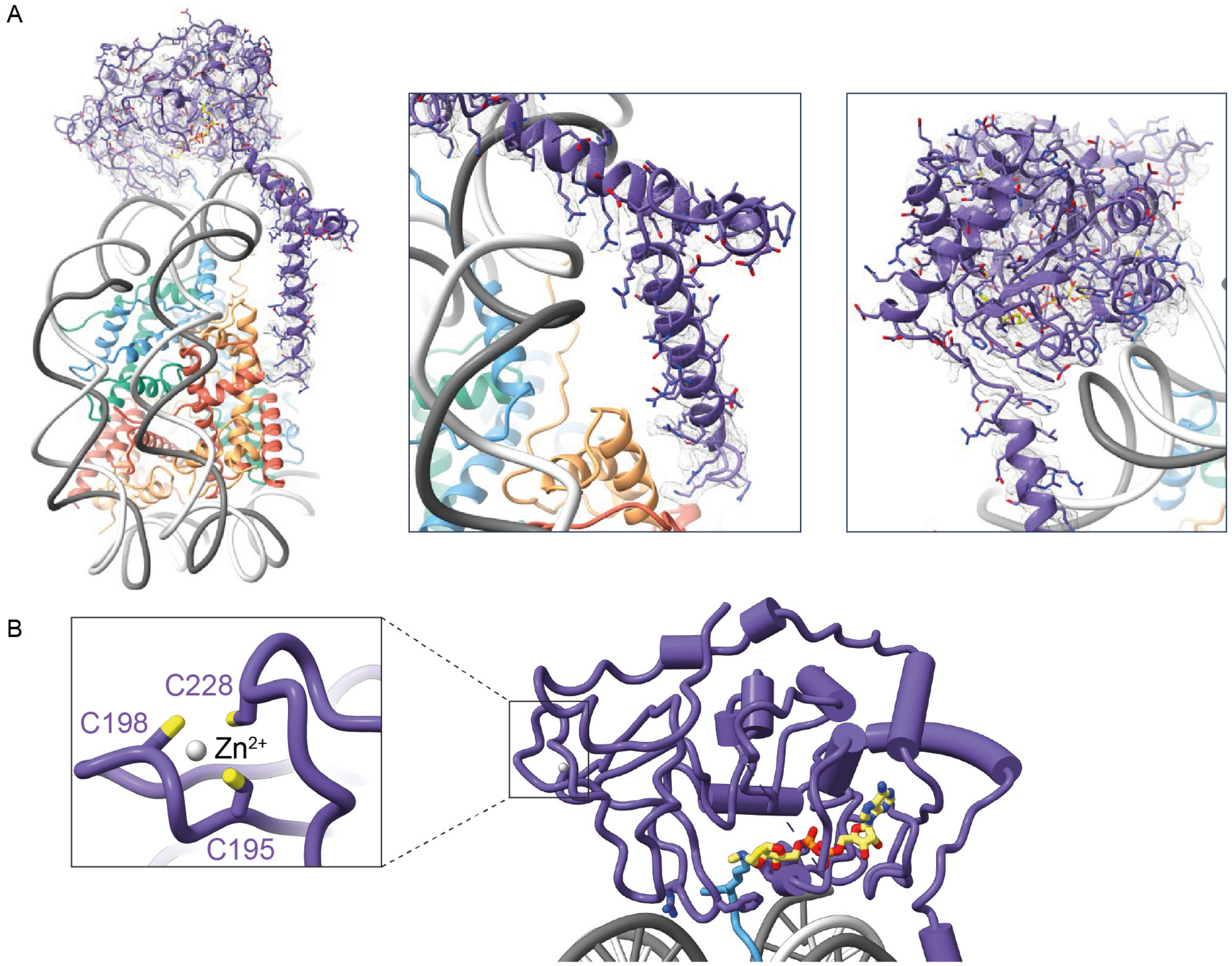

**Supplementary Fig. 5. A**, Overall and zoom-in views of the cryo-EM map of structural elements of SIRT7 in the cryo-EM map (mesh representation) of the SIRT7:H3K36MTU-nucleosome complex. **B**, Detail of the $Zn^{2+}$-binding residues of the SIRT7 catalytic domain.

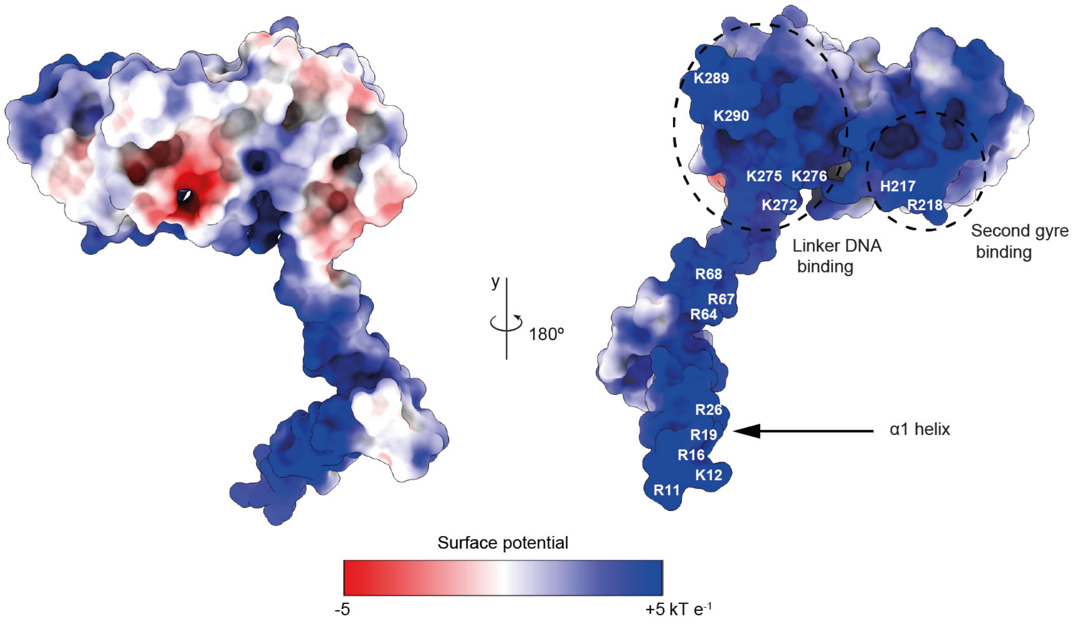

**Supplementary Fig. 6.** Surface potential of SIRT7 with highlighted nucleosome-binding regions.

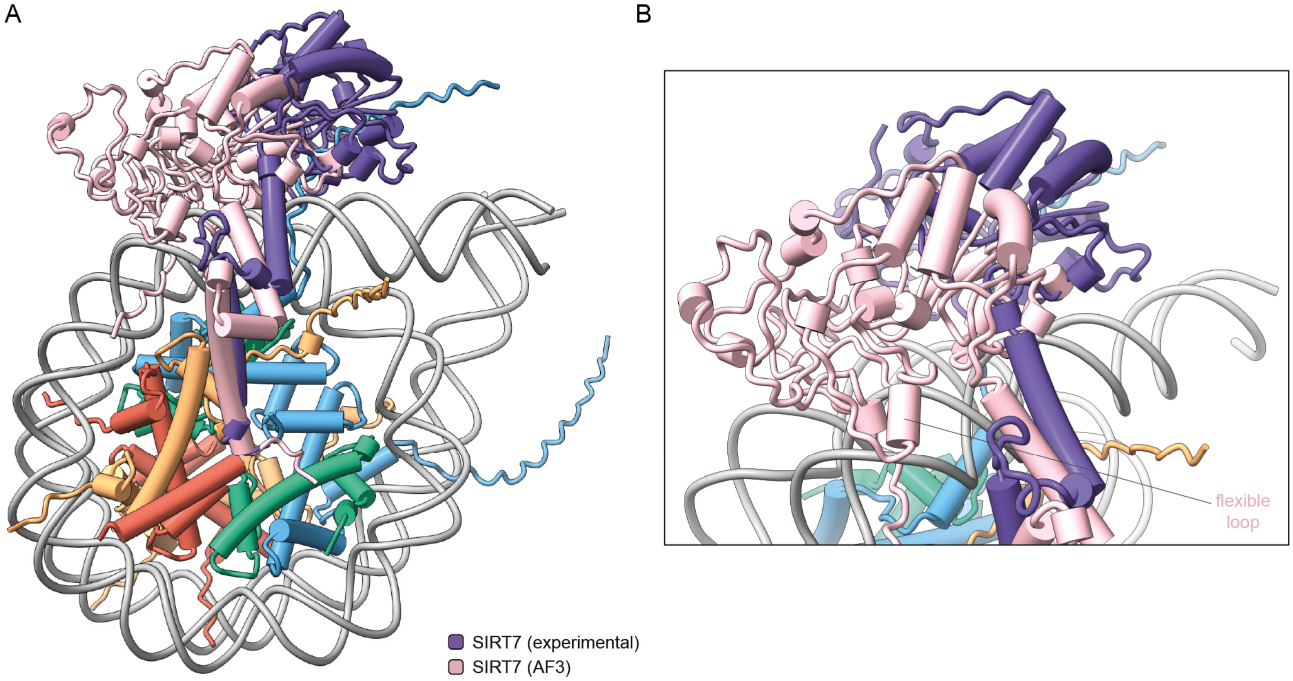

**Supplementary Fig. 7. AlphaFold 3 prediction of the SIRT7:nucleosome complex. A**, Overlay of the AF3 SIRT7:nucleosome complex and our experimental model of SIRT7 bound to the H3K36MTU nucleosome (not shown). Structures were aligned by the histone octamer. **B**, Detail of the flexible loop within the catalytic domain (aa 120–138) and its AF3-predicted interaction with DNA. This region has lower confidence than the rest of the catalytic domain according to AF3.

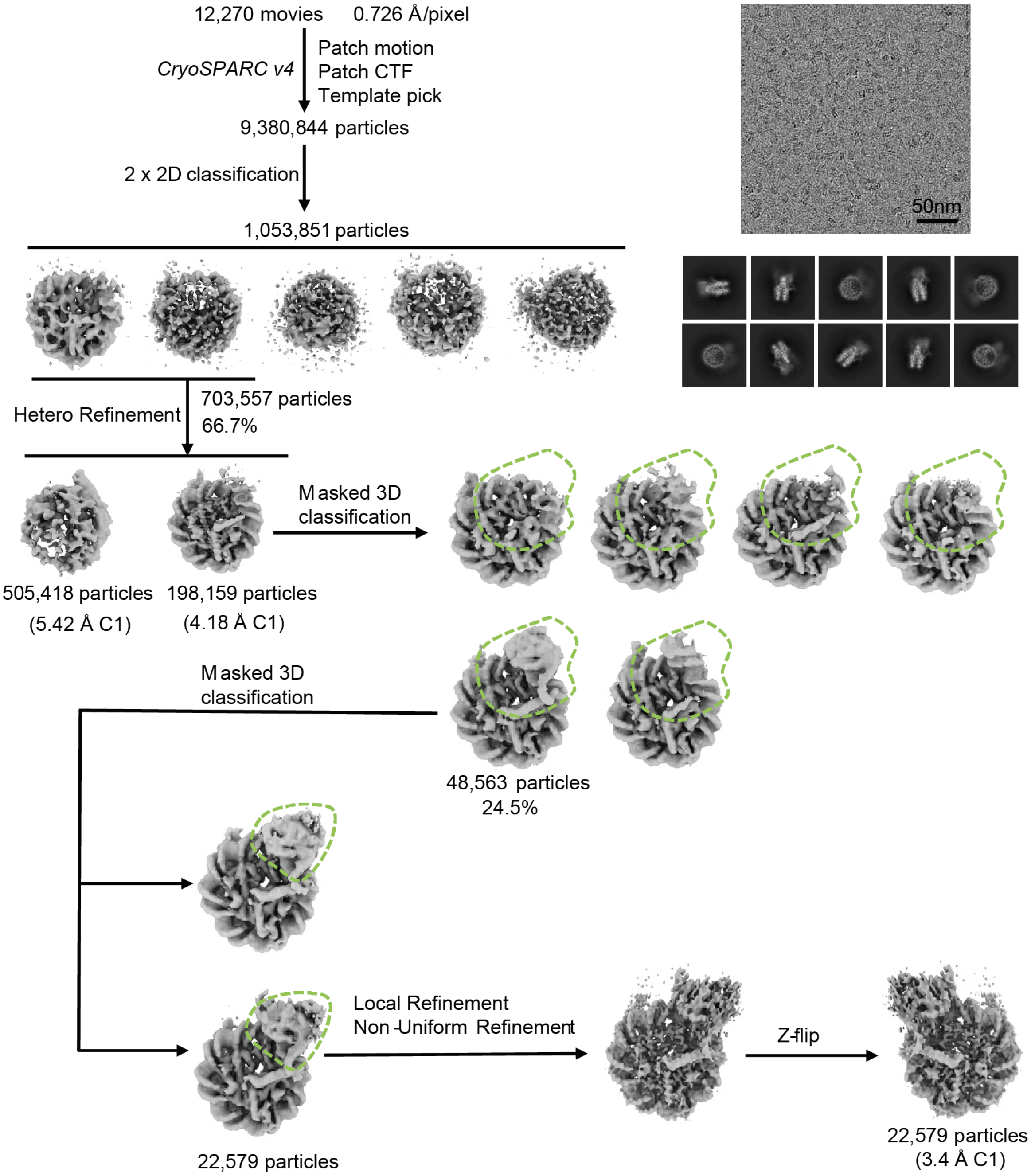

**Supplementary Fig. 8. SIRT7:H3K18DTU complex data processing workflow.** Processing workflow for SIRT7:H3K18DTU complex showing 2D class averages, 3D classification and refinement steps. Focused classification with the regions covered by the mask are also shown.

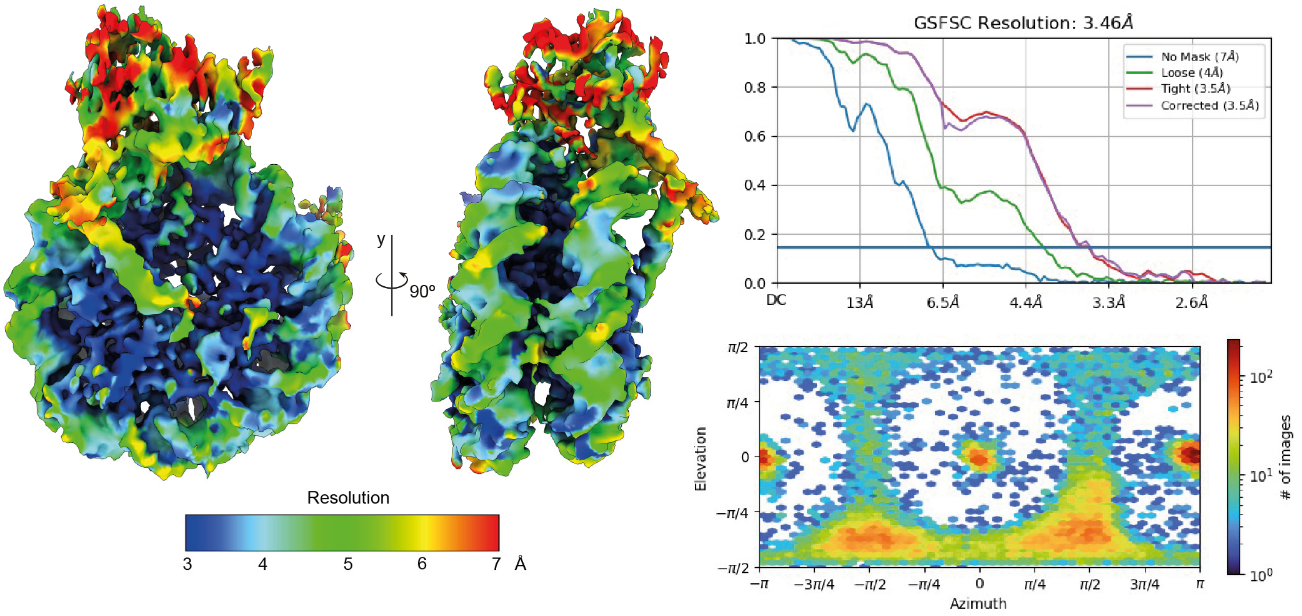

**Supplementary Fig. 9.** Local resolution map, Fourier Shell Correlation (FSC) curve and particle distribution orientation of the SIRT7:H3K18DTU-nucleosome complex.

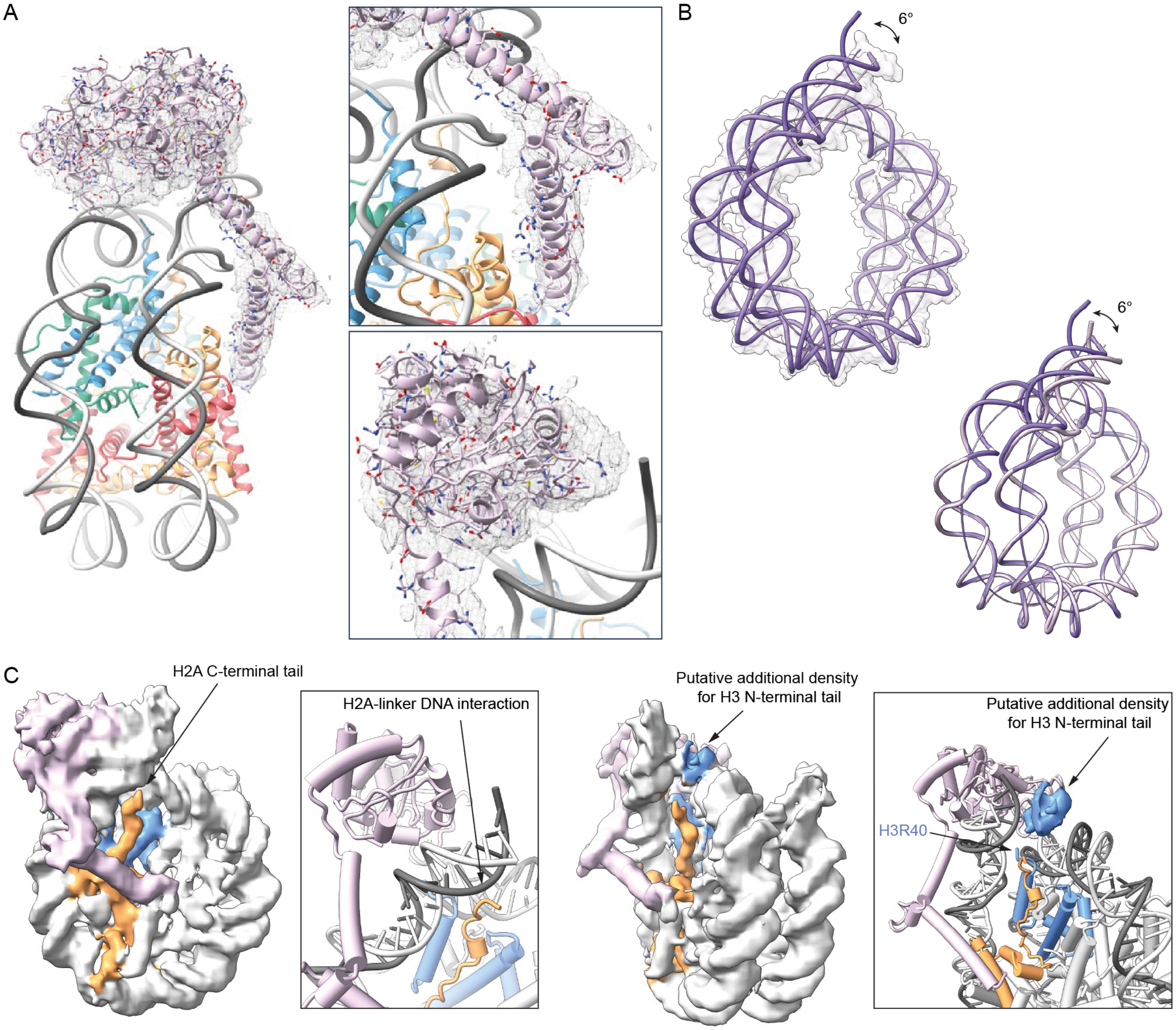

**Supplementary Fig. 10. A**, Overall and zoom-in views of the cryo-EM map of structural elements of SIRT7 in the cryo-EM map (mesh representation) of the SIRT7:H3K18DTU-nucleosome complex. **B**, Measurement of the relative bending of the linker DNA axes, between the H3K36- (purple) and H3K18 (light pink)-bound structures. The H3K18-bound model is shown either as surface (top) or cartoon (bottom). **C**, H2A C-terminal and H3 N-terminal densities suggesting histone:DNA interactions that may further explain the relative bending of linker DNA.

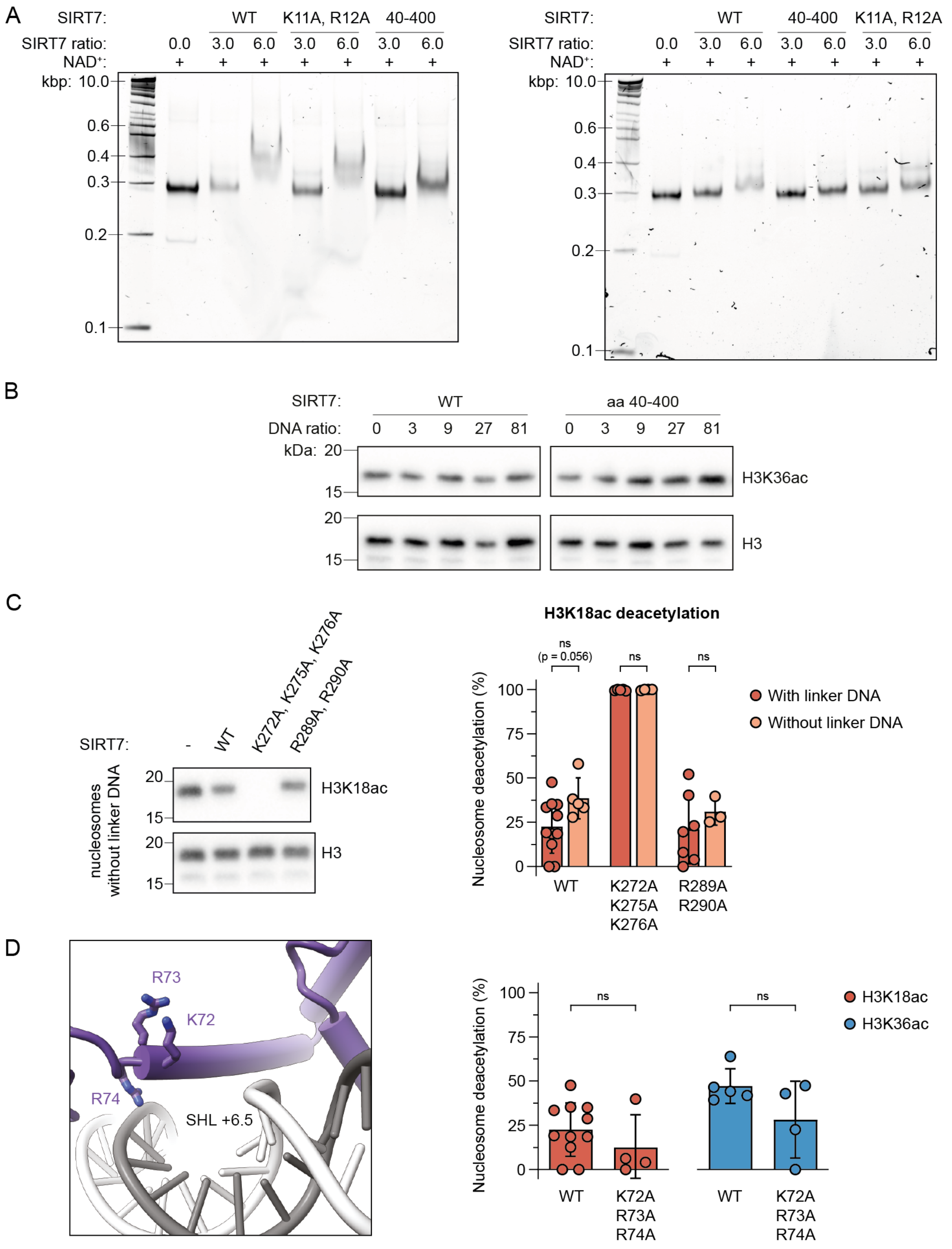

**Supplementary Fig. 11. Additional data on SIRT7 mutant activity. A**, EMSA replicates of unmodified nucleosomes (187 bp DNA) with wild type SIRT7, truncated SIRT7 (aa 40-400) and the K11A, R12A mutant. **B**, Inhibitory effect of free DNA (187 bp) on SIRT7 activity, measured by western blot. **C**, Activity of wild type SIRT7 and selected mutants on H3K18ac-modified nucleosomes with standard DNA (187 bp) and 601 DNA without linker ends (147 bp). Error bars represent mean ± SD ($n \geq 3$), ns: $p > 0.05$. **D**, Detailed interactions of the third N-terminal helix and quantification of nucleosome deacetylation normalized to H3 loading, at a concentration of wild type SIRT7 or the K72A, R73A and R74A mutant of 50 nM for H3K18ac and 3 nM for H3K36ac experiments. These mutations were observed to further promote the activation by 5S RNA[1]. Nucleosome concentration: 200 nM. Error bars represent mean ± SD ($n \geq 4$), ns: $p > 0.05$.

### Supplementary Tables

**Supplementary Table 1. Cryo-EM data collection, refinement and validation statistics for SIRT7-nucleosome complexes.**

| Data collection and processing | SIRT7:H3K36MTU nucleosome complex (EMD-XXXX) (PDB XXXX) | SIRT7:H3K18DTU nucleosome complex (EMD-XXXX) (PDB XXXX) |
|---|---|---|
| Electron Microscope | Titan Krios G4 (ColdFEG, Falcon4i) | Titan Krios G4 (ColdFEG, Falcon4i, SelectrisX) |
| Nominal Magnification | 96kx | 165kx |
| Voltage (kV) | 300 | 300 |
| Recorded Micrographs | 8 623 | 12 270 |
| Electron exposure (e–/Å2) | 50 | 50 |
| Defocus range (μm) | 0.8-2.5 | 0.8-2.5 |
| Pixel size (Å) | 0.82 | 0.726 |
| Symmetry imposed | C1 | C1 |
| Initial particle images (no.) | 7 266 736 | 9 149 090 |
| Final particle images (no.) | 337 476 | 22 579 |
| Map resolution (Å) | 2.8 | 3.5 |
| FSC threshold | 0.143 | 0.143 |
| Map resolution range (Å) | 5.0-2.0 | 7.0-3.0 |
| Map sharpening B factor (Å2) | -76.1 | -42.9 |
| **Refinement** | | |
| Initial model used (PDB code) | 3LZ0 | 3LZ0 |
| Model composition<br>Atoms (hydrogens)<br>Protein residues<br>Nucleotide residues<br>Ligands | 14789 (24)<br>1086<br>298<br>ZSL: 1<br>ZN: 1 | 14643 (0)<br>1081<br>296 |
| *B* factors ($Å^2$)<br>Protein<br>Nucleotide<br>Ligand | 0.00/87.72/36.16<br>11.59/155.09/70.70<br>0.50/97.91/2.94 | 232.91/799.07/391.79<br>257.47/587.40/348.43 |
| R.m.s. deviations<br>Bond lengths (Å)<br>Bond angles (°) | 0.004 (0)<br>0.569 (2) | 0.004 (0)<br>0.739 (31) |
| Validation<br>MolProbity score<br>Clashscore<br>Poor rotamers (%) | 1.71<br>14.67<br>2.09 | 1.83<br>21.52<br>0.33 |
| Ramachandran plot<br>Favored (%)<br>Allowed (%)<br>Disallowed (%) | 97.47<br>2.44<br>0.09 | 98.02<br>1.98<br>0.00 |

### Chemical synthesis

*General methods*

All commercial reagents and solvents were of analytical grade and used without further purification. $H_2O$ was of MilliQ grade unless otherwise stated and obtained from a Thermo Scientific GenPure UF system. Reactions were monitored by HPLC-MS analysis using a Shimadzu MS2020 instrument equipped with a Waters Acquity UPLC C18 column (for peptide analysis) and a Waters Acquity UPLC C4 column (for protein analysis), and UV diode array and single quadrupole analysis systems. Gradients of eluent I (0.05% HCOOH in $H_2O$) and eluent II (0.05% HCOOH in MeCN) were used as mobile phase. Purification and chromatographic analyses were performed by reverse-phase HPLC on Agilent 1260 systems, with Zorbax 300SB-C18 columns #1 (7 µm, 250×21.2 mm, 300 Å, for preparative purification), #2 (5 µm, 250×9.4 mm, 300 Å, for semi-preparative purification), and #3 (5 µm, 150×4.60 mm, 300 Å, for analysis), using gradients of eluent III (0.1% TFA in $H_2O$) and eluent IV (0.1% TFA in MeCN/$H_2O$ 90:10) at a flow rate of 20 mL/min (preparative purification), 4 mL/min (semi-preparative purification) or 1 mL/min (analysis). Identification of purification fractions and of the final products consisted on high-resolution MS analysis using a Waters Xevo G2-XS instrument equipped with UV diode array and quadrupole-time-of-flight analysis systems, by direct injection.

*Peptide synthesis*

Peptide synthesis was performed based on the protocol by Guidotti *et al.*, 2020[2].

To prepare hydrazide-modified resins, 2-chlorotrityl chloride resin (0.5 g, 1.55 mmol/g, Merck, cat. # 8.55017) was swelled in a round-bottom flask with DMF (3 mL) for 15 min at room temperature, followed by cooling in an ice bath and addition of a solution of hydrazine (1 mL, 1.65 M hydrazine monohydrate and 2.45 M *i*$Pr_2$EtN in DMF) dropwise under gentle stirring. The mixture was allowed to reach room temperature and further stirred for 1 h, after which MeOH (0.1 mL) was added and the mixture was stirred for additional 10 min. The resin was then transferred to vessels for manual solid phase peptide synthesis (SPPS), washed with DMF (3 × 2 min), and dried under suction. A solution of Fmoc-Val-OH or Fmoc-Ser(*t*Bu)-OH (2.5 equiv.), 1-[bis(dimethylamino)methylene]-1*H*-1,2,3-triazolo[4,5-*b*]pyridinium 3-oxid hexafluorophosphate (HATU, 2.38 equiv.) and *i*$Pr_2$EtN (5.0 equiv.) was prepared in DMF (for a 0.5 M concentration of HATU), incubated for 1 min at room temperature, and added to the dried resin. The reaction vessel was agitated gently for 1 h at room temperature, after which the resin was dried under suction and washed with DMF (3 × 2 min), $CH_2Cl_2$ (3 × 2 min) and MeOH (3 × 2 min), and dried under vacuum overnight. Loading of the resin was determined as reported[3]. Briefly, 1-3 mg resin aliquots were incubated with piperidine/DMF (4 mL, 20:80, v/v) for 30 min at room temperature, and the absorption of the solution at 301 nm was used to calculate the loading (0.6-0.7 mmol/g, Supplementary Fig. 12).

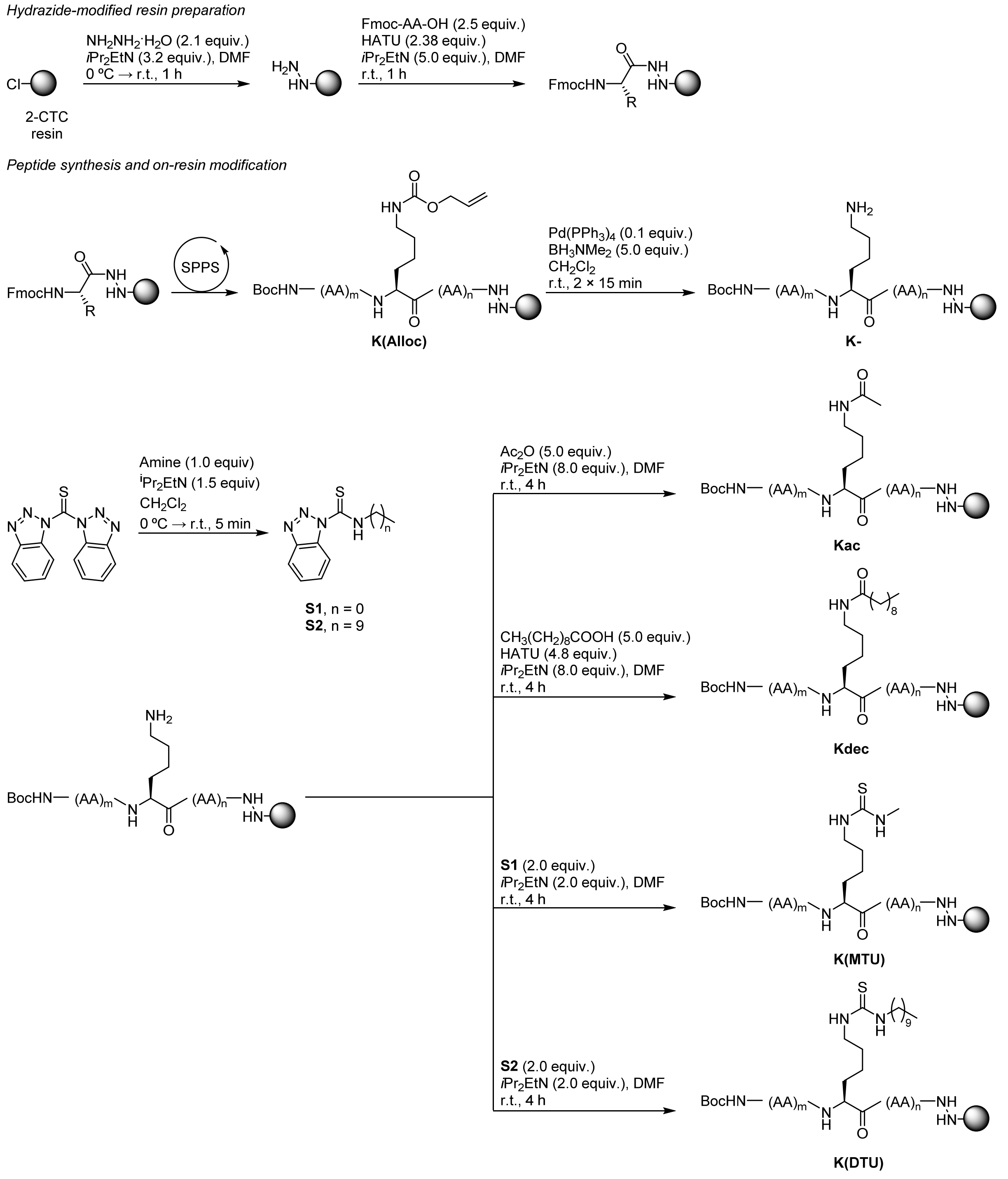

**Supplementary Fig. 12.** Summary of peptide synthesis and on-resin modification methods.

Standard Fmoc/t-Bu SPPS was performed on a Tribute peptide synthesizer (Gyros Protein Technologies, Sweden) using hydrazide-modified resins (0.05 or 0.1 mmol), *O*-(1*H*-6-chlorobenzotriazole-1-yl)-1,1,3,3-tetramethyluronium hexafluorophosphate (HCTU) as coupling reagent, DMF as solvent, and protected amino acids as follow: Boc-Ala-OH, Boc-Thz-OH, Fmoc-Ala-OH, Fmoc-Arg(Pbf)-OH, Fmoc-Gln(Trt)-OH, Fmoc-Gly-OH, Fmoc-

His(Trt)-OH, Fmoc-Leu-OH, Fmoc-Lys(Alloc)-OH, Fmoc-Lys(Boc)-OH, Fmoc-Pro-OH, Fmoc-Ser(*t*Bu)-OH, Fmoc-Thr(*t*Bu)-OH, Fmoc-Tyr(*t*Bu)-OH and Fmoc-Val-OH. The following pseudo-proline building blocks were used to improve synthetic yields: Fmoc-Ala-Thr[ψ(Me,Me)pro]-OH, Fmoc-Gln(Trt)-Thr[ψ(Me,Me)pro]-OH and Fmoc-Lys(Boc)-Ser[ψ(Me,Me)pro]-OH (see peptide sequences below). Fmoc deprotection was performed using piperidine/DMF (20:80, v/v) for 2 × 3 min. Amino acids (5.0 equiv.) were pre-activated with HCTU (4.8 equiv.) and *i*$Pr_2EtN$ (7.0 equiv.) at a concentration of 0.25 M, added to the resin, and incubated by shaking for 30 min at room temperature. Reactions were repeated when indicated (double coupling, see peptide sequences below). Resin washes were performed with DMF. Resin aliquots were cleaved using a mixture of TFA/TIPS/$H_2O$ (95:2.5:2.5) for reaction monitoring by HPLC-MS.

Peptide sequences prepared (underlined residues were double-coupled, and residues in italics correspond to pseudo-proline building blocks):

**K18Alloc:** Boc-A R T K *QT* A R *KS* T G G K A P R K(Alloc) Q L *AT* K A A R K S-NHNH-resin

**K36Alloc:** Boc-Thz P A T G G V K(Alloc) K P H R Y R P G T V-NHNH-resin

*On-resin lysine modification*

On-resin selective deprotection of K(Alloc) was performed in syringe reactors with a mixture $Pd(PPh_3)_4$ (0.1 equiv.) and $BH_3NMe_2$ (5.0 equiv.) in $CH_2Cl_2$ for 15 min at room temperature. The resins were then washed with $CH_2Cl_2$ (3 × 2 min) and the treatment was repeated. Resins were finally washed with MeOH (3 × 2 min) and dried under vacuum for storage, or used directly in the next step (Supplementary Fig. 12).

On-resin formation of Kac and Kdec was achieved by incubation of Alloc-deprotected peptidyl-resin samples (1.0 equiv.) with a solution of $Ac_2O$ (5.0 equiv.) and *i*$Pr_2EtN$ (8.0 equiv.), or decanoic acid (5.0 equiv.), HATU (4.8 equiv.) and *i*$Pr_2EtN$ (8.0 equiv.), in DMF (at a concentration of acylating reagent of 0.1 M) for 4 h at room temperature, followed by washes with DMF (3 × 2 min), $CH_2Cl_2$ (3 × 2 min) and MeOH (3 × 2 min), and drying under vacuum.

On-resin thiourea formation was achieved as reported[4]. A solution of $MeNH_2$ (2 M in DMF, 2.0 equiv.) and *i*$Pr_2EtN$ (3.0 equiv.), or n-decylamine (2.0 equiv.) and *i*$Pr_2EtN$ (3.0 equiv.), in anhydrous $CH_2Cl_2$ (to a concentration of amine of 40 mM) was added dropwise to a solution of bis(benzotriazol-1-yl)methanethione (75 mM in anhydrous $CH_2Cl_2$, 2.0 equiv.) in an ice bath over 5 min. The resulting mixtures were concentrated under reduced pressure, diluted with a solution of *i*$Pr_2EtN$ (75 mM in anhydrous DMF, 2.0 equiv.), and quickly added to the Alloc-deprotected peptidyl-resin (1.0 equiv.). The reactions were shaken for 4 h at room temperature, followed by washes with DMF (3 × 2 min), $CH_2Cl_2$ (3 × 2 min) and MeOH (3 × 2 min), and drying under vacuum.

All reactions were followed by test cleavage of peptidyl-resin aliquots using a mixture of TFA/TIPS/$H_2O$ (95:2.5:2.5) and monitoring by HPLC-MS, and repeated when necessary.

*Peptide cleavage and deprotection*

Overall peptide deprotection and cleavage from resin was achieved by shaking of peptidyl-resin samples with a mixture of TFA/2,2′-(ethylenedioxy)diethanethiol (DODT)/TIPS/$H_2O$ (92.5:2.5:2.5:2.5, 1 mL per 100 mg of resin) for 2-3 h at room temperature. The resulting solutions were concentrated under $N_2$ stream and triturated with ice-cold $Et_2O$. The mixtures were centrifuged (3000 g, 3 min, 4 ºC), and the pellets were washed again with ice-cold $Et_2O$, separated by centrifugation (3000 g, 3 min, 4 ºC), dried under $N_2$ stream and purified by preparative HPLC as described.

The following peptides were obtained (Supplementary Fig. 13):

| Peptide | Modification | Purity* | Formula | HRMS | Calcd. mass | Synth. yield |
|---|---|---|---|---|---|---|
| **S3** | K18- | 99% | $C_{125}H_{232}N_{50}O_{36}$ | 3009.761 | 3009.786 | 20% |
| **S4** | K18ac | >90% | $C_{127}H_{234}N_{50}O_{37}$ | 3051.800 | 3051.797 | 6% |
| **S5** | K18dec | 90% | $C_{135}H_{250}N_{50}O_{37}$ | 3163.920 | 3163.922 | 6% |
| **S6** | K18MTU | 97% | $C_{127}H_{235}N_{51}O_{36}S$ | 3082.758 | 3082.785 | 3% |
| **S7** | K18DTU | 99% | $C_{136}H_{253}N_{51}O_{36}S$ | 3208.900 | 3208.926 | 4% |
| **S8** | K36ac | 94% | $C_{87}H_{142}N_{30}O_{22}S$ | 1991.065 | 1991.064 | 30% |
| **S9** | K36dec | 84% | $C_{95}H_{158}N_{30}O_{22}S$ | 2103.194 | 2103.189 | 22% |
| **S10** | K36MTU | 83% | $C_{87}H_{143}N_{31}O_{21}S_2$ | 2021.722 | 2022.052 | 16% |
| **S11** | K36DTU | 92% | $C_{96}H_{161}N_{31}O_{21}S_2$ | 2147.841 | 2148.192 | 23% |

*Peptide purity measured by integration of HPLC chromatograms at 214 nm. See Supporting Fig. 18 for HPLC traces.

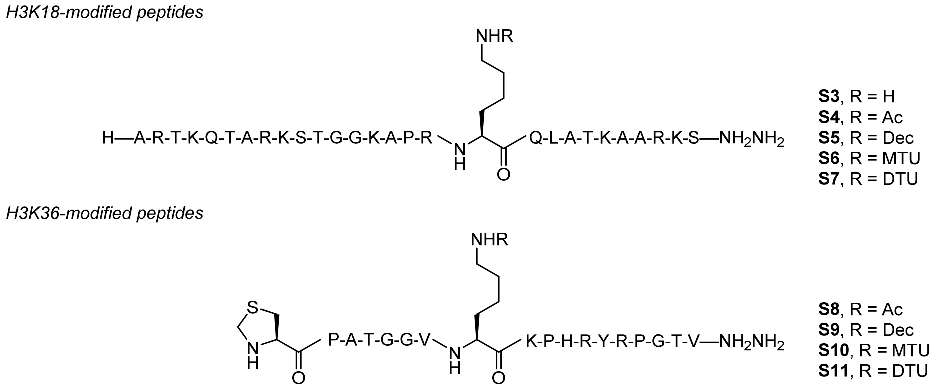

**Supplementary Fig. 13.** Chemical structure of synthesized peptides.

*Semi-synthesis of modified full-length histone H3*

Histone H3 semi-synthesis was modified from the protocol by Guidotti *et al.*, 2020[2].

Peptide hydrazides (3.0 equiv.) were converted into peptide thioesters by *in situ* pyrazole formation, as reported[5]. Lyophilized peptides were dissolved in a dispersion of 4-mercaptophenylacetic acid (MPAA, 60 equiv.) in acidic ligation buffer (6 M guanidinium chloride, 0.2 M $NaH_2PO_4$, 0.2 M MPAA, pH 3) to a concentration of 10 mM, followed by addition of acetylacetone (0.5 M in $H_2O$, 7.5 equiv.) and stirring at room temperature for 2-4 h under Ar atmosphere. The reaction was followed by HPLC-MS and, when full conversion to the MPAA thioester was detected, the crude mixtures were stored at -20 ºC for ligation within the following days (Supplementary Fig. 14).

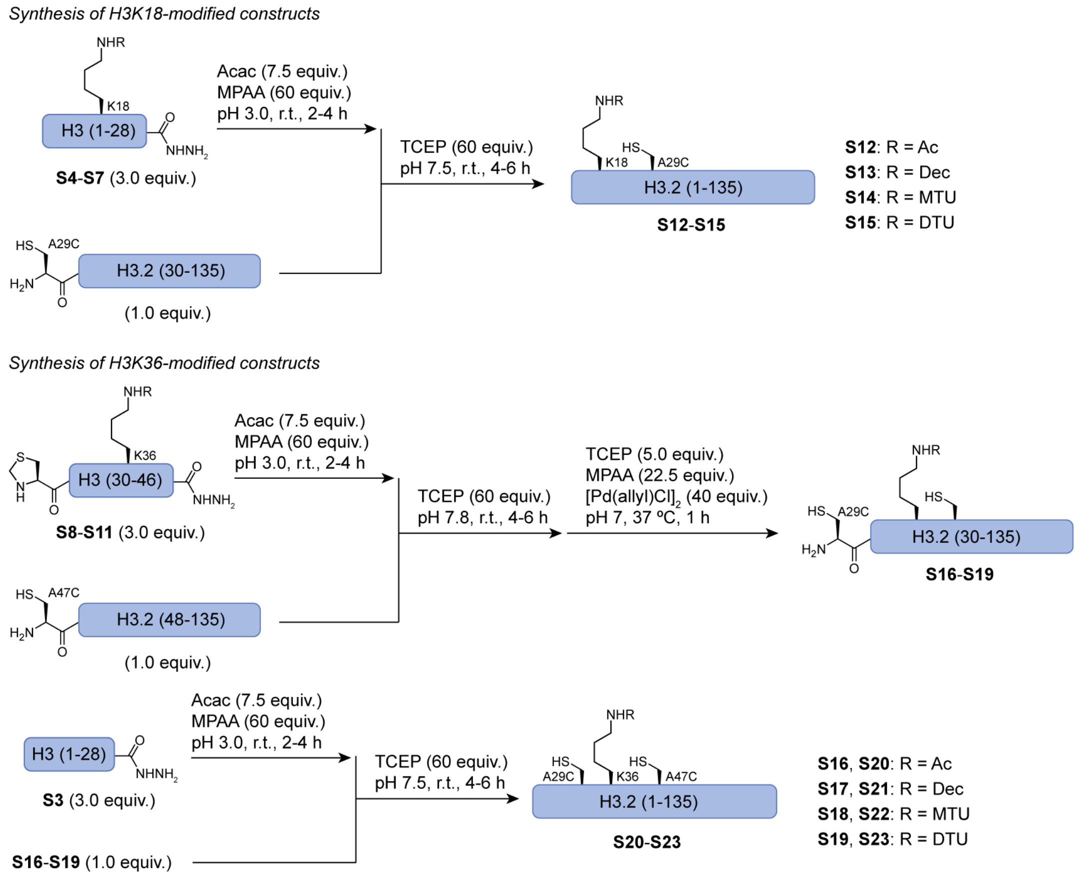

**Supplementary Fig. 14.** Summary of histone semi-synthesis methods. Acac: acetylacetone.

Peptide thioesters were linked to N-terminal Cys-containing histone fragments (see recombinant expression and purification below) by native chemical ligation. The histone fragment lyophilized powder (1.0 equiv.) was dissolved in the peptide thioester crude mixture (3.0 equiv.) by vortexing, and the concentration of histone fragment was adjusted to ~1 mM with ligation buffer (6 M guanidinium chloride, 0.2 M $NaH_2PO_4$, pH 7). Then, tris(2-carboxyethyl)phosphine hydrochloride (TCEP, 0.5 M in ligation buffer, 60 equiv.) was added, the pH was adjusted carefully to 7.8 (for ligation to H3(47-135) fragments) or 7.5 (for ligation to H3(29-135) fragments) using NaOH (5 M or 1 M in $H_2O$), and the mixture was allowed to stir at room temperature for 4-6 h under Ar atmosphere. Reactions were followed by HPLC-MS and analytical HPLC and, either diluted with eluents III and IV and purified directly by semi-preparative HPLC for full-length constructs, or treated for thiazolidine deprotection in the case of ligation to H3(47-135) fragments (Supplementary Fig. 14).

Thiazolidine deprotection was achieved by addition of TCEP (0.5 M in ligation buffer, 5.0 equiv.), MPAA (0.2 M in ligation buffer, 22.5 equiv.) and $[Pd(allyl)Cl]_2$ (0.1 M in ligation buffer, 40 equiv.) to the crude ligation mixture, and vigorous shaking at 37 ºC for 1 h under Ar atmosphere (Supplementary Fig. 14). Thereafter, DTT (0.5 M in $H_2O$) was added to final concentration of 250 mM, the mixture was vortexed extensively, and it was further diluted to half with $H_2O$/MeCN 1:1 before centrifugation (21130 g, 10 min, 25 ºC). The pellet was washed twice with DTT (250 mM in $H_2O$/MeCN 3:1 with 0.05% TFA), and the combined supernatant was diluted with eluents III and IV and purified by semi-preparative HPLC.

The following full-length histones were obtained:

| Protein | Modification | Purity* | Formula | HRMS* | Calcd. MW | Synth. yield |
|---|---|---|---|---|---|---|
| **S12** | K18ac | >85% | $C_{671}H_{1131}N_{215}O_{187}S_3$ | 15299.6 | 15299.0 | 34% |
| **S13** | K18dec | >90% | $C_{679}H_{1147}N_{215}O_{187}S_3$ | 15411.2 | 15411.1 | 36% |
| **S14** | K18MTU | >95% | $C_{671}H_{1132}N_{216}O_{186}S_4$ | 15330.0 | 15330.0 | 42% |
| **S15** | K18DTU | >95% | $C_{680}H_{1150}N_{216}O_{186}S_4$ | 15456.2 | 15456.2 | 42% |
| **S20** | K36ac | 97% | $C_{671}H_{1131}N_{215}O_{187}S_4$ | 15330.8 | 15331.0 | 12% |
| **S21** | K36dec | 92% | $C_{679}H_{1147}N_{215}O_{187}S_4$ | 15443.0 | 15443.2 | 19% |
| **S22** | K36MTU | >80% | $C_{671}H_{1132}N_{216}O_{186}S_5$ | 15362.0 | 15362.1 | 22% |
| **S23** | K36DTU | >85% | $C_{680}H_{1150}N_{216}O_{186}S_5$ | 15488.0 | 15488.3 | 13% |

*Protein purity measured by integration of HPLC chromatograms at 214 nm. HRMS calculated by molecular weight deconvolution with MassLynx MaxEnt1. See Supporting Figs. 19, 22, 23 for HPLC traces and mass spectra.

### Histone expression and purification

#### *Expression and purification of H3(29-135) and H3(47-135) constructs*

H3.2 C110A fragment expression and purification was performed based on the protocol by Guidotti *et al.*, 2020[2].

*E. coli* BL21 (DE3) cells were transformed with a pET30 plasmid encoding for the corresponding truncated histone H3.2 with an N-terminal A→C mutation, an N-terminal fusion of 6xH-SUMO, and the mutation C110A common in H3 protein semi-synthesis. Cells were cultured in 5 L LB medium with 50 µg/mL kanamycin at 37 ºC with shaking at 200 rpm until $OD_{600}$ = 0.6-0.8. Then, 1 M isopropyl β-D-1-thiogalactopyranoside (IPTG) was added to a concentration of 150 mM and the cells were cultured for additional 4 h at 37 ºC. Cell pellets were obtained by centrifugation (4000 g, 20 min, 4 ºC), suspended in H3 lysis buffer (60 mL, 20 mM Tris-HCl, 200 mM NaCl, 1 mM EDTA, 1 mM 2-mercaptoethanol, pH 7.5) supplemented with protease inhibitors (cOmplete EDTA free, Merck), and flash-frozen for storage at -70 ºC. Thereafter, the suspension was thawed and lysed by sonication for 10 min (15 s on, 45 s off pulses). The insoluble fraction was collected by centrifugation (15000 g, 30 min, 4 ºC), washed with standard H3 lysis buffer (2 × 60 mL), with H3 lysis buffer supplemented with 1% Triton X-100 (1 × 60 mL), and with standard H3 lysis buffer (1 × 60 mL). The resulting pellet was suspended in a solubilization buffer (6 M guanidinium chloride, 50 mM Tris-HCl, 100 mM NaCl, 5 mM imidazole, 1 mM 2-mercaptoethanol, pH 7.5) by vigorous shaking and stirring for 2-4 h at 4 ºC, centrifuged (15000 g, 30 min, 4 ºC), and the supernatant was purified in several batches by Ni-NTA FPLC. The column was washed with buffer E (6 M urea, 150 mM Tris-HCl, 150 mM NaCl, 50 mM imidazole, pH 7.5), and the sample was eluted by a mixture 60:40 (v/v) of buffer E and buffer F (6 M urea, 150 mM Tris-

HCl, 150 mM NaCl, 500 mM imidazole, pH 7.5). Fractions containing the corresponding 6x-SUMO-H3 construct were identified by SDS-PAGE (15% acrylamide gel), combined, mixed with commercial Ulp1 SUMO protease (Merck, cat. # SAE0067) and dialyzed against 2 L of a cleavage buffer (1 M urea, 75 mM Tris-HCl, 150 mM NaCl, 5 mM 1,4-dithiothreitol (DTT), 25 mM L-arginine, pH 7.5). SUMO cleavage was verified by HPLC-MS, and the sample was dialyzed into 1% AcOH and lyophilized. The resulting powder was either treated with $[Pd(allyl)Cl]_2$, MPAA and TCEP (in the case of thiazolidine formation due to formaldehyde contamination) and purified, or purified directly by preparative HPLC. Typical yields were of 2.5-5 mg of lyophilized protein powder per liter of bacterial culture.

6xH-SUMO-H3(29-135) A29C, C110A sequence (red: removed by Ulp1 cleavage)

MGSSHHHHHH GSGLVPRGSA SMSDSEVNQE AKPEVKPEVK PETHINLKVS DGSSEIFFKI KKTTPLRRLM EAFAKRQGKE MDSLRFLYDG IRIQADQTPE DLDMEDNDII EAHREQIGG**C** PATGGVKKPH RYRPGTVALR EIRRYQKSTE LLIRKLPFQR LVREIAQDFK TDLRFQSSAV MALQEASEAY LVGLFEDTNL AAIHAKRVTI MPKDIQLARR IRGERA

6xH-SUMO-H3(47-135) A47C, C110A sequence (red: sequence after Ulp1 cleavage)

MGSSHHHHHH GSGLVPRGSA SMSDSEVNQE AKPEVKPEVK PETHINLKVS DGSSEIFFKI KKTTPLRRLM EAFAKRQGKE MDSLRFLYDG IRIQADQTPE DLDMEDNDII EAHREQIGG**C** LREIRRYQKS TELLIRKLPF QRLVREIAQD FKTDLRFQSS AVMALQEASE AYLVGLFEDT NLAAIHAKRV TIMPKDIQLA RRIRGERA

*Expression and purification of wild type H2A, H2B, H3 and H4*

Wild type H2A, H2B, and H4, and H3 C110A, were produced based on literature procedures[6].

Transformed *E.coli* BL21 (DE3) cells were cultured in 6 L LB medium with 100 µg/mL ampicillin at 37 ºC with shaking at 200 rpm until $OD_{600}$ of ~0.6-0.8. Then, 1 M IPTG was added to a final concentration of 1 mM, and the cells were cultured for additional 3-4 h at 37 ºC. Cells were harvested, lysed, and the pellets were prepared as mentioned before for 6xH-SUMO-H3 constructs. Pellets were then resuspended in a solubilization buffer (6 M guanidinium chloride, 20 mM Tris-HCl, 1 mM EDTA, 1 mM 2-mercaptoethanol, pH 7.5) by stirring at 4 ºC overnight, centrifuged (15000 g, 30 min, 4 ºC), and the supernatant was dialyzed against 2 L of a urea-containing buffer (7 M urea, 10 mM Tris-HCl, 100 mM NaCl, 1 mM EDTA, 5 mM 2-mercaptoethanol, 0.2 mM PMSF, pH 7.5) and centrifuged again (15000 g, 30 min, 4 ºC). The new supernatant was purified by IEX FPLC using a 0→100% linear gradient of NaCl between the previous urea-containing buffer (100 mM NaCl) and high-salt version of the same buffer (1500 mM NaCl). Fractions containing the desired histone were identified by SDS-PAGE (15% acrylamide gel), combined, and purified by preparative HPLC. Typical yields were of 7-25 mg of lyophilized protein powder per liter of bacterial culture.

Wild type H2A sequence (*H. sapiens* H2A type 2-A):

SGRGKQGGKA RAKAKSRSSR AGLQFPVGRV HRLLRKGNYA ERVGAGAPVY MAAVLEYLTA EILELAGNAA RDNKKTRIIP RHLQLAIRND EELNKLLGKV TIAQGGVLPN IQAVLLPKKT ESHHKAKGK

Wild type H2B sequence (*H. sapiens* H2B type 1-K):

PEPAKSAPAP KKGSKKAVTK AQKKDGKKRK RSRKESYSVY VYKVLKQVHP DTGISSKAMG IMNSFVNDIF ERIAGEASRL AHYNKRSTIT SREIQTAVRL LLPGELAKHA VSEGTKAVTK YTSAK

Wild type H3 sequence (*H. sapiens* H3.2 with C110A mutation):

ARTKQTARKS TGGKAPRKQL ATKAARKSAP ATGGVKKPHR YRPGTVALRE IRRYQKSTEL LIRKLPFQRL VREIAQDFKT DLRFQSSAVM ALQEASEAYL VGLFEDTNLA AIHAKRVTIM PKDIQLARRI RGERA

Wild type H4 sequence (*H. sapiens* H4):

SGRGKGGKGL GKGGAKRHRK VLRDNIQGIT KPAIRRLARR GGVKRISGLI YEETRGVLKV FLENVIRDAV TYTEHAKRKT VTAMDVVYAL KRQGRTLYGF GG

### Gel electrophoresis of protein, DNA and nucleosome samples

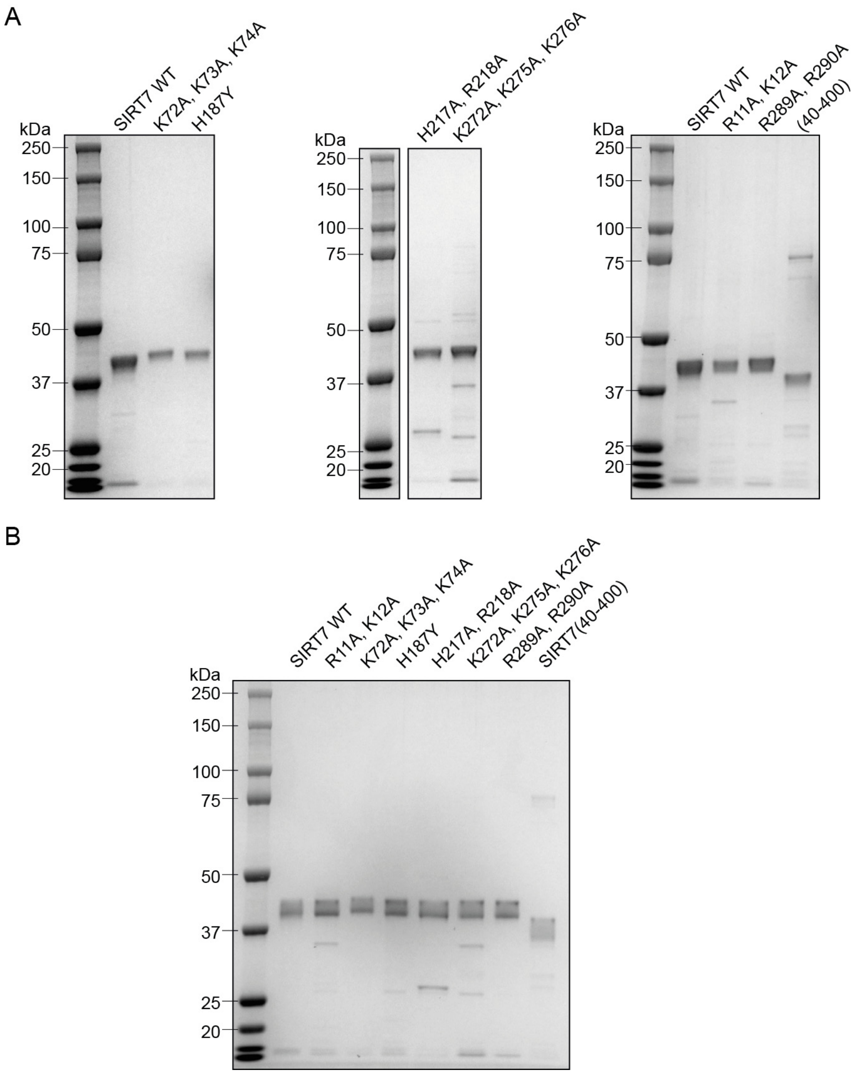

**Supplementary Fig. 15.** SDS-PAGE analysis of recombinant SIRT7 preparations employed in this work. **a**, Example of SDS-PAGE analysis of each of the constructs stained with Coomasie Blue, as used for SIRT7 quantification. **b**, Summary gel containing ~0.3 µg of each SIRT7 construct, stained with Coomasie Blue. See Supporting Figs. 20–21 for mass spectra.

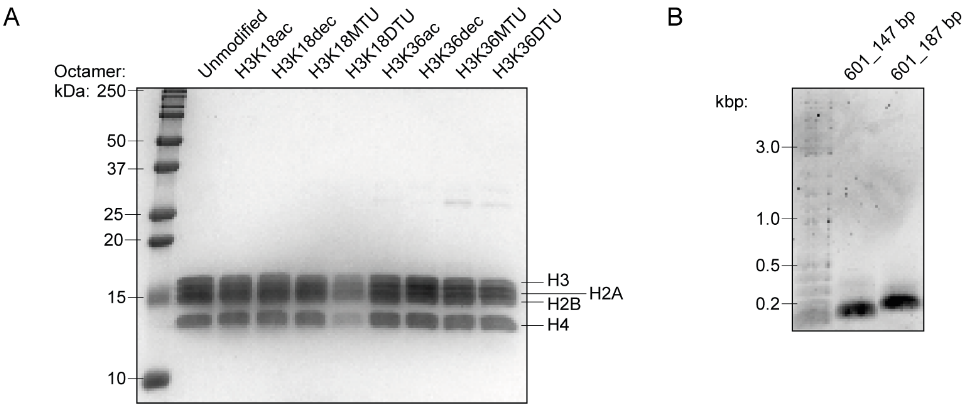

**Supplementary Fig. 16. A**, SDS-PAGE analysis of all histone octamers employed in this work. Octamers are labelled according to H3 modifications, as they all contain wild type H2A, H2B and H4. Unmodified octamers contain H3.2 C110A. **B**, Agarose gel electrophoretic analysis of DNA samples, stained with GelRed.

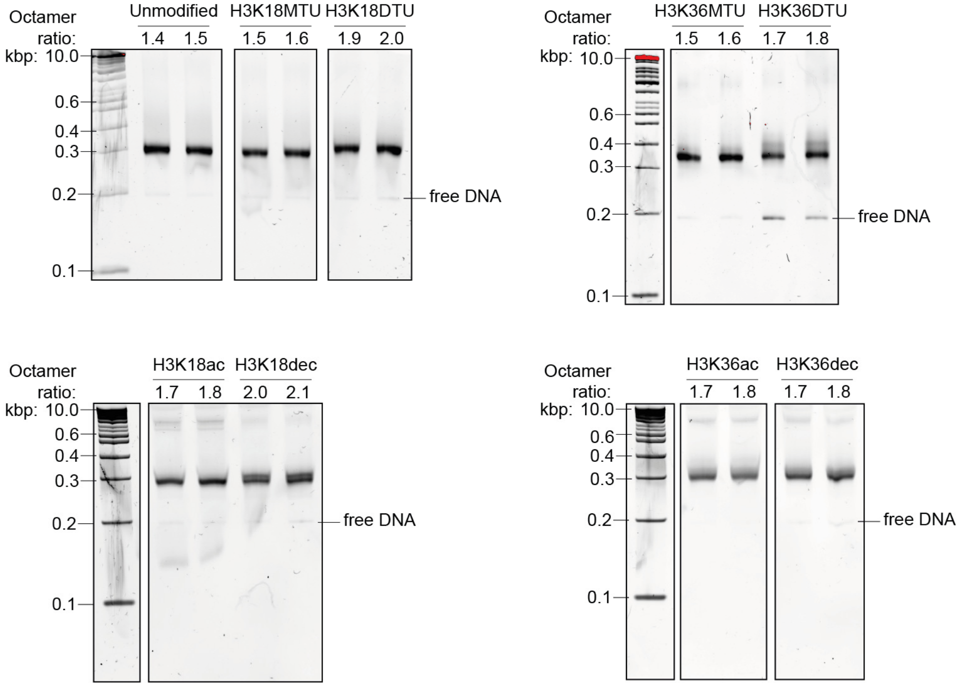

**Supplementary Fig. 17.** Examples of native gel electrophoretic analysis of nucleosome samples, stained with GelRed.

### Purity tracesB

*Peptide HPLC traces*

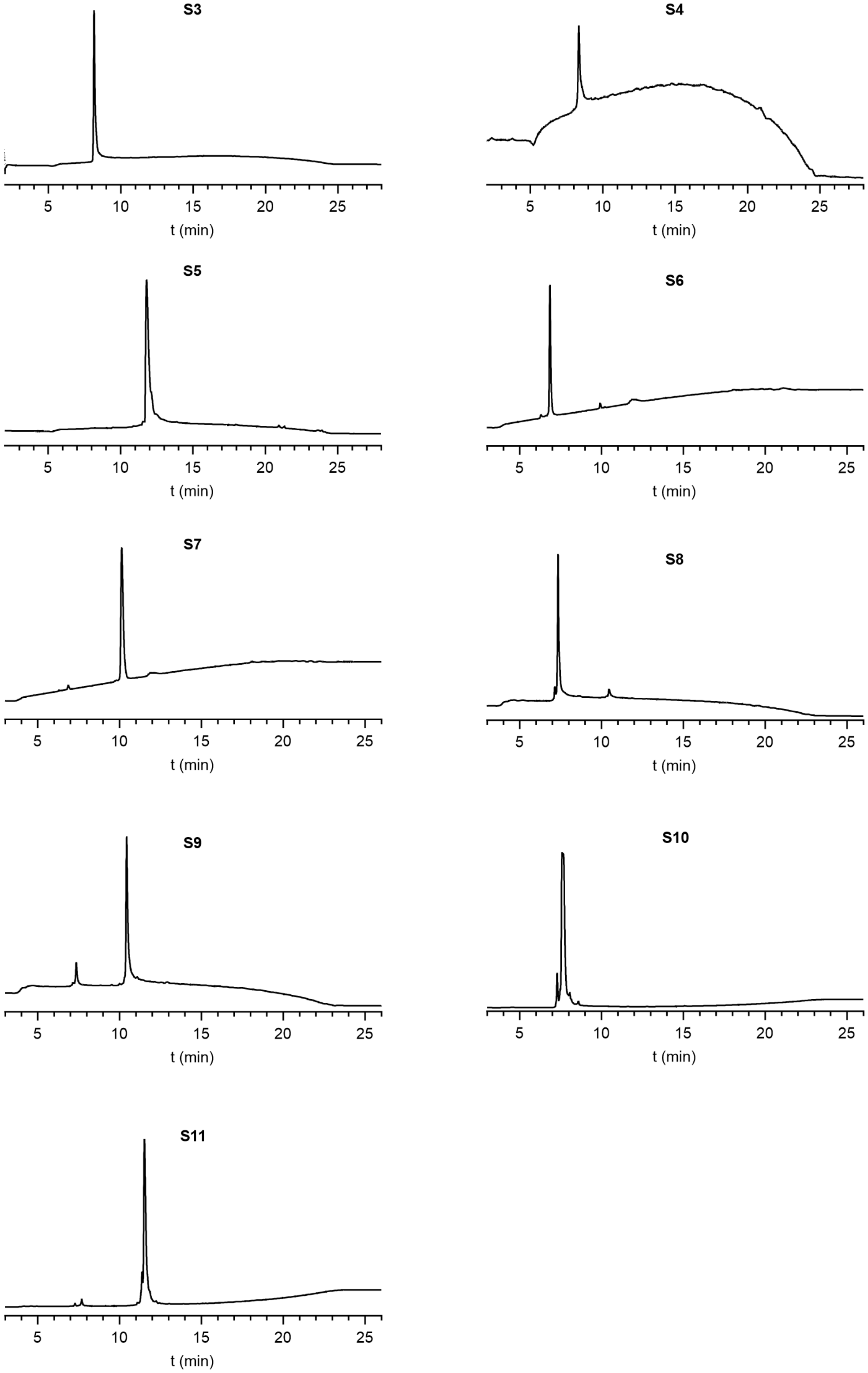

**Supplementary Fig. 18.**

*Full-length modified histone H3 HPLC traces*

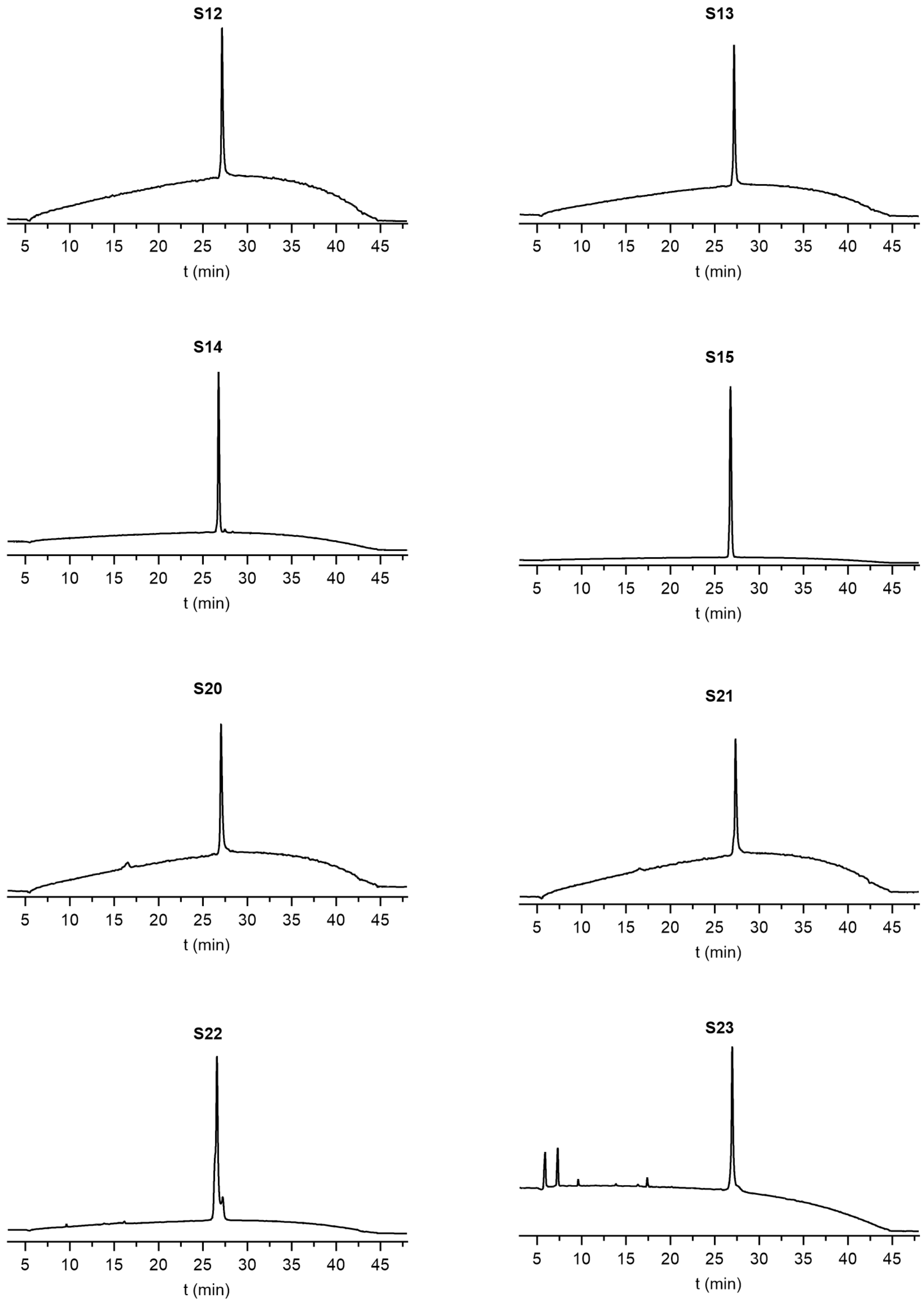

**Supplementary Fig. 19**. Note: compound **S23** was injected dissolved in unfolding buffer, which gave extra signals between 5 and 8 min.

### Mass spectra

*SIRT7 mass spectra (left) and deconvoluted mass (right)*

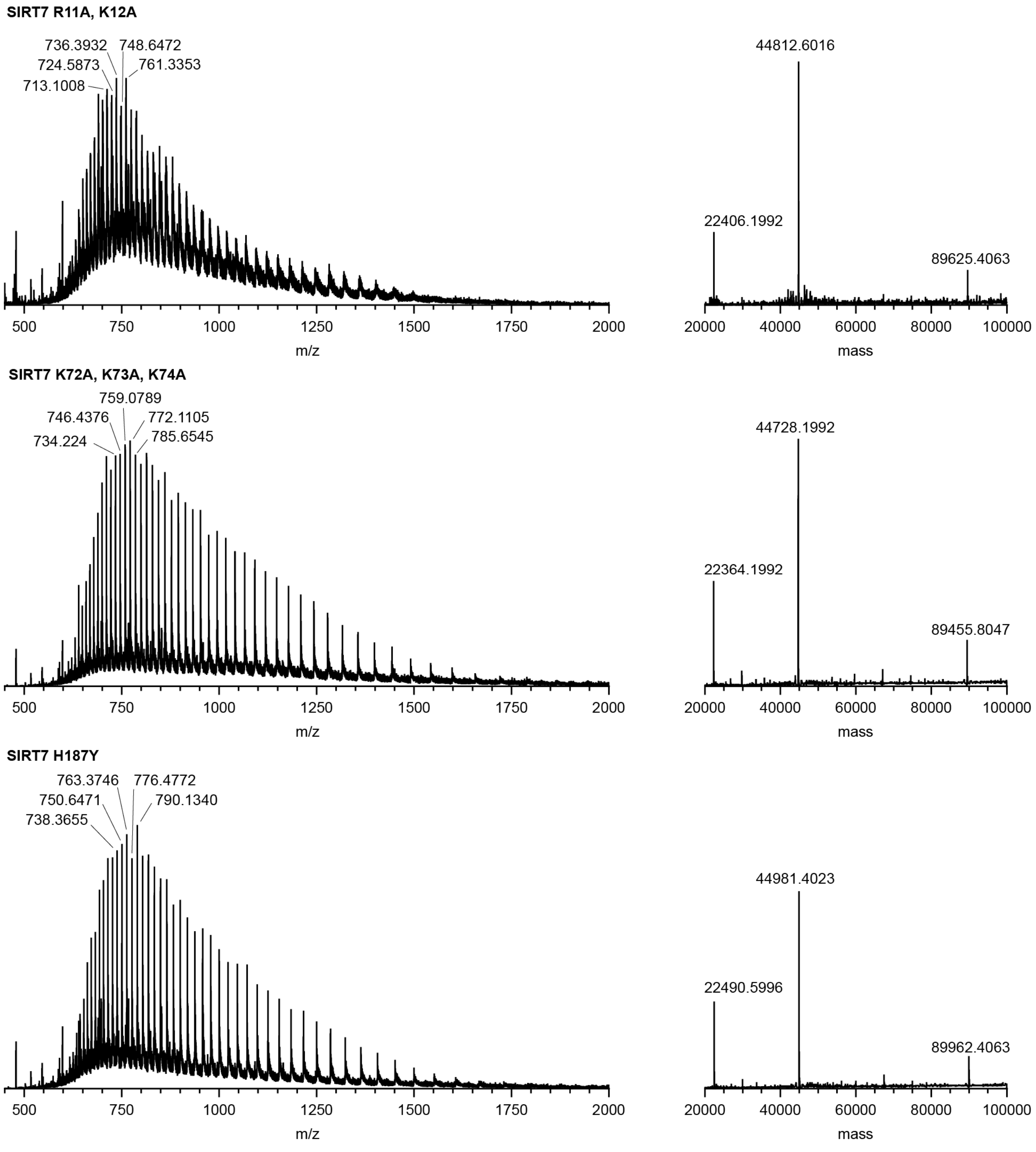

**Supplementary Fig. 20.**

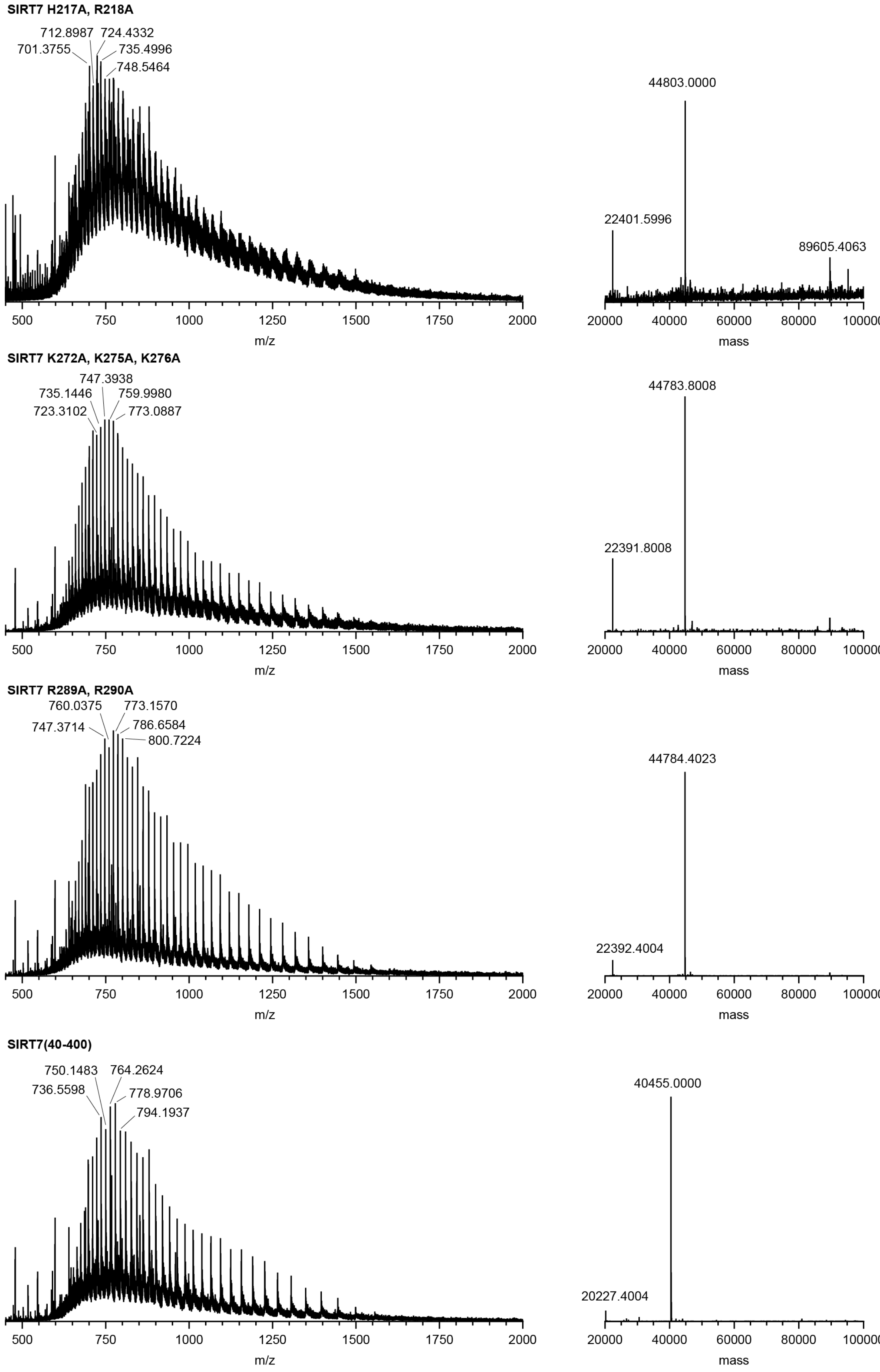

**Supplementary Fig. 21.**

*Full-length modified histone H3 mass spectra (left) and deconvoluted mass (right)*

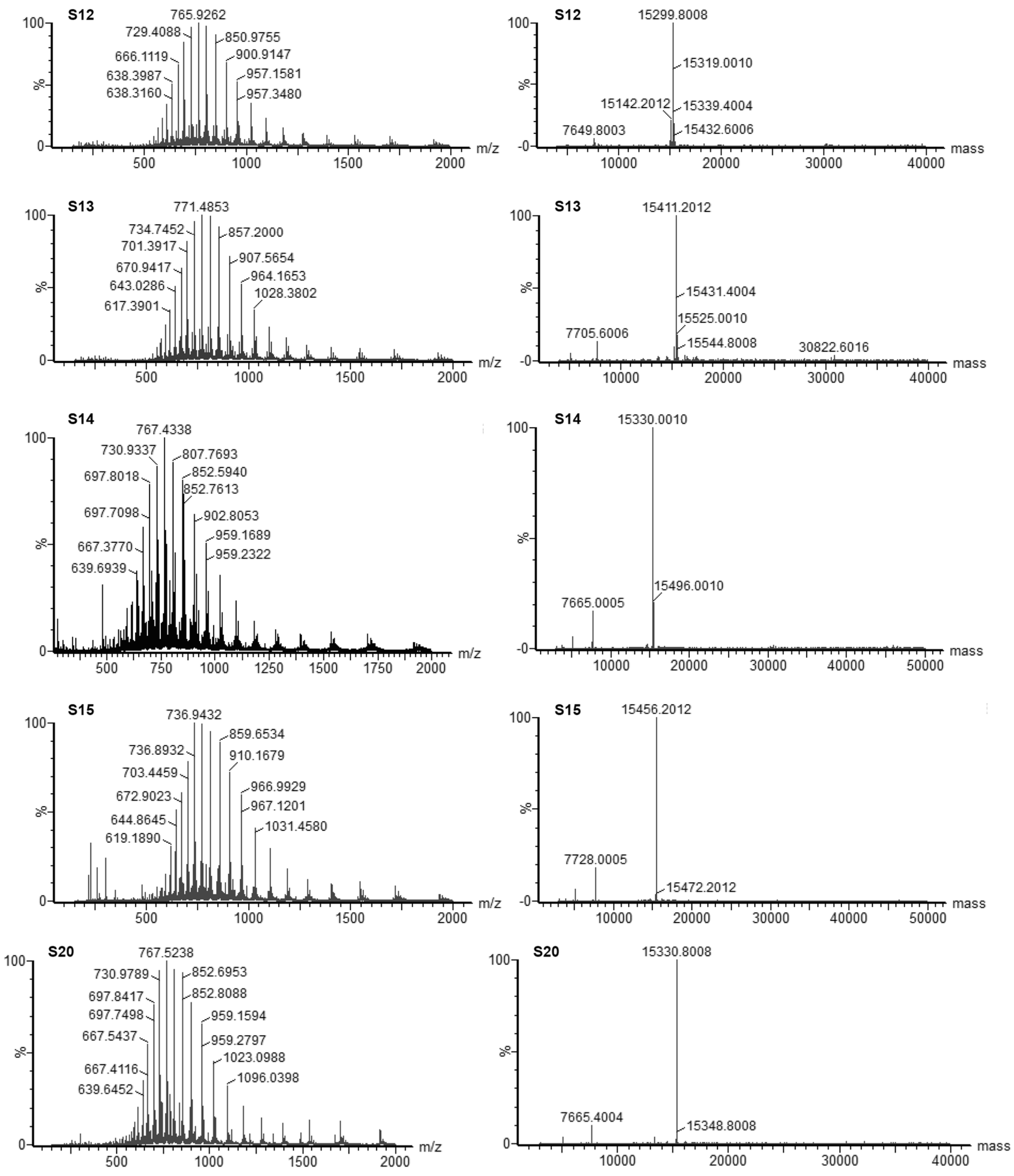

**Supplementary Fig. 22.**

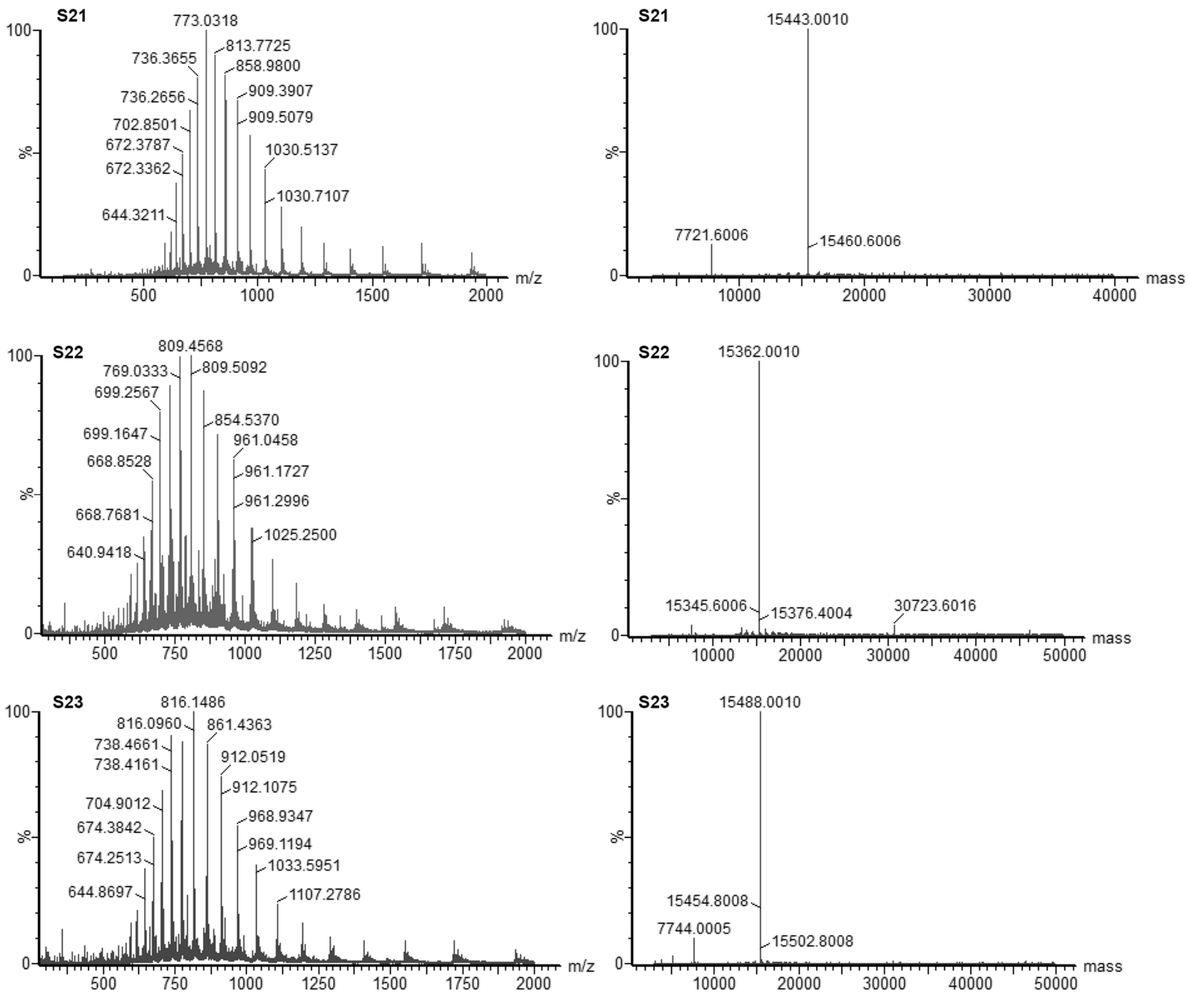

**Supplementary Fig. 23.**

### Supporting references

1. Tong, Z. et al. SIRT7 Is an RNA-Activated Protein Lysine Deacylase. *ACS Chem Biol* **12**, 300-310 (2017).
2. Guidotti, N. & Fierz, B. Semisynthesis and Reconstitution of Nucleosomes Carrying Asymmetric Histone Modifications. *Methods Mol Biol* **2133**, 263-291 (2020).
3. Eissler, S. et al. Substitution determination of Fmoc-substituted resins at different wavelengths. *J Pept Sci* **23**, 757-762 (2017).
4. Troelsen, K.S. et al. Mitochondria-targeted inhibitors of the human SIRT3 lysine deacetylase. *RSC Chem Biol* **2**, 627-635 (2021).
5. Flood, D.T. et al. Leveraging the Knorr Pyrazole Synthesis for the Facile Generation of Thioester Surrogates for use in Native Chemical Ligation. *Angew Chem Int Ed Engl* **57**, 11634-11639 (2018).
6. Dyer, P.N. et al. Reconstitution of nucleosome core particles from recombinant histones and DNA. *Methods Enzymol* **375**, 23-44 (2004).
